# Dynamic Multiplexed Control and Modeling of Optogenetic Systems Using the High-Throughput Optogenetic Platform, Lustro

**DOI:** 10.1101/2023.12.19.572411

**Authors:** Zachary P. Harmer, Jaron C. Thompson, David L. Cole, Victor M. Zavala, Megan N. McClean

## Abstract

The ability to control cellular processes using optogenetics is inducer-limited, with most optogenetic systems responding to blue light. To address this limitation we leverage an integrated framework combining Lustro, a powerful high-throughput optogenetics platform, and machine learning tools to enable multiplexed control over blue light-sensitive optogenetic systems. Specifically, we identify light induction conditions for sequential activation as well as preferential activation and switching between pairs of light-sensitive spit transcription factors in the budding yeast, *Saccharomyces cerevisiae*. We use the high-throughput data generated from Lustro to build a Bayesian optimization framework that incorporates data-driven learning, uncertainty quantification, and experimental design to enable the prediction of system behavior and the identification of optimal conditions for multiplexed control. This work lays the foundation for designing more advanced synthetic biological circuits incorporating optogenetics, where multiple circuit components can be controlled using designer light induction programs, with broad implications for biotechnology and bioengineering.

**Graphical abstract:** 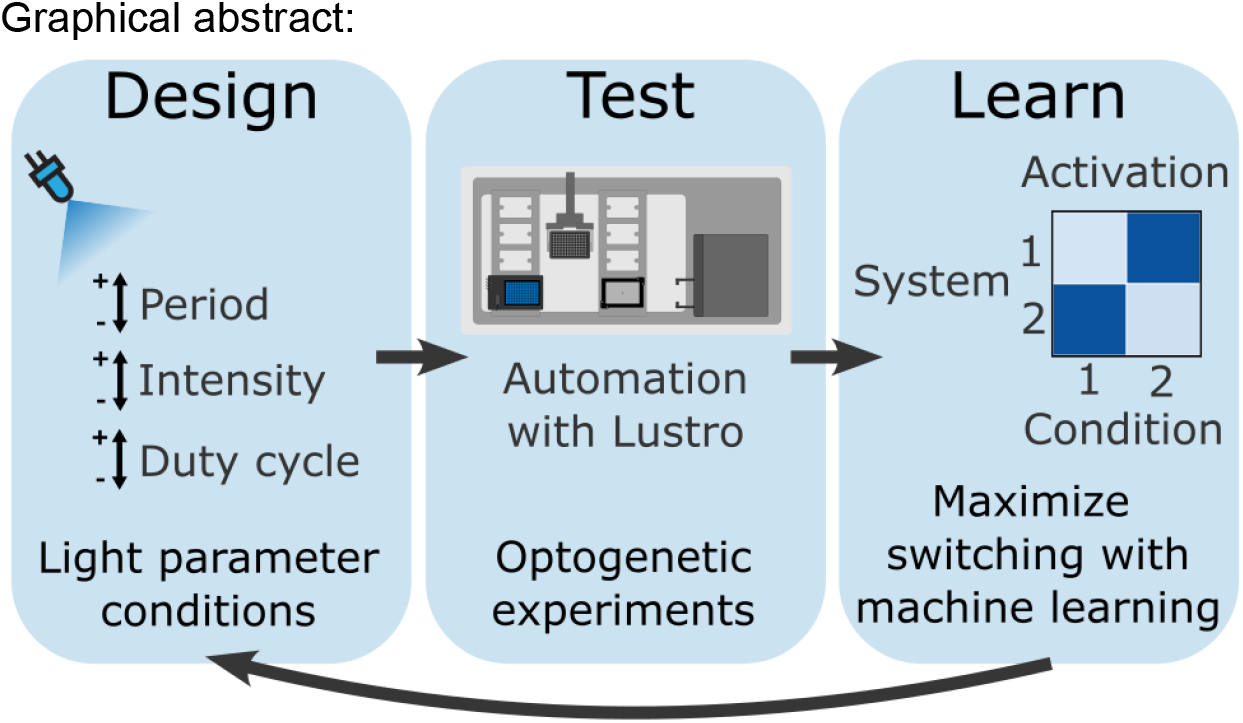

## Introduction

Optogenetics leverages genetically encoded light-sensitive proteins fused to biological effectors to precisely control cellular behavior in response to light^1–3^. Such optogenetic technologies empower researchers to orchestrate cellular processes with exquisite spatiotemporal precision. Optogenetic technologies have found diverse applications, ranging from modulating gene expression^4–7^, dissecting intricate signaling pathways^8–10^, manipulating protein localization^11–15^, or inducing targeted protein degradation^16,17^. If independent and simultaneous control of distinct optogenetic systems is possible, engineering multiple optogenetic systems into the same cell or community of cells allows for a higher degree of control over complex biological functions. This independent control can be achieved via multiplexing, where multiple control signals are sent over a shared medium, in this case light^18,20^. One approach is orthogonal multiplexing, where optogenetic systems responsive to different wavelengths of light are used. However, a significant limitation in optogenetics is the fact that most protein photoswitches are responsive to blue light^18^. One approach to overcome this limitation is dynamic multiplexing, where specific light induction programs of the same wavelength, but with different duration and period of illumination pulses, are used to selectively activate optogenetic systems.

Benzinger and Khammash developed one strategy for dynamic multiplexed control of optogenetic systems by taking advantage of EL222 mutants with different response kinetics, that is different activation and reversion timescales in the light and dark, respectively^20^. The authors built a falling edge detector from two different EL222 mutants, and this circuit generated a distinct response profile to light induction programs relative to another optogenetic system based on the cryptochome CRY2 and its binding partner CIB1. With sufficiently differentiated response kinetics, multiplexed control of optogenetic systems is possible without the need for such additional circuitry. However, the response kinetics of optogenetics systems *in vivo* are not well understood and are difficult to measure at high throughput. The ability to rapidly construct and characterize optogenetic systems, coupled with data-driven modeling, presents a promising avenue to navigate this challenge, cutting down the search space to find maximally differentiated outputs for tailored multiplexing schemes.

In this work, we present a strategy for taking advantage of the native differences in response kinetics between protein photoswitches by using a previously described automated high-throughput optogenetic platform, Lustro^7,19^, to identify light induction programs that allow for dynamic multiplexed control over optogenetic systems. We use the high-throughput measurement capabilities of Lustro to characterize a set of 13 blue light-responsive optogenetic split transcription factors (TFs). Optogenetic split TFs use complementary dimerization domains fused to a DNA-binding domain (DBD) and an activation domain (AD), such that light-induced dimerization of the protein pair reconstitutes the split TF and expression of the gene of interest is induced. We selected optogenetic split TFs for developing multiplexing strategies as their activity can be readily measured using a fluorescent protein reporter, control of gene expression is useful for a broad range of biological applications, and many mutants of optical dimerizers with different response kinetics are known (see Table 1). We used this high-throughput characterization to empirically identify specific sets of light induction programs that result in distinct activation levels for different blue light-sensitive optogenetic systems, allowing us to multiplex control over them using those light induction programs. We identified conditions for sequential activation, where differences in light sensitivity between optogenetic systems result in differential activation of each optogenetic system. We also identified conditions for “switching,” where one light induction program preferentially activates one optogenetic system over a second, but switching to a second light program results in preferential activation of the second optogenetic system over the first. We combine the high-throughput characterization with a Bayesian machine learning framework that aims to predict and optimize objectives for optogenetic control. Furthermore, we highlight the powerful synergy between high-throughput data collection and predictive models, showcasing how their integration can unravel the complexities of optogenetic systems, paving the way for a new era of finer cellular control and optimization.

**Table 1.** Optogenetic split TFs characterized in this work (Figure 1). The eMagA/eMagB (and variants) dimerizer pair was engineered into a heterodimerizer from VVD (a homodimerizer taken from *Neurospora crassa*) and used to create two-component split TFs^23,24^. The CRY2/CIB1 (and variants) dimerizer pair was derived from *Arabidopsis thaliana* and engineered into a two-component split TF^25^. EL222 and its variants are a homodimerizer taken from *Erythrobacter litoralis* and engineered into a single-component split transcription factor (with the VP16AD) for use in eukaryotes^26^. Split TFs derived from heterodimerizers (CRY2/CIB1, eMagA/eMagB, and their variants) use Gal4AD and Gal4DBD^7,28^. Additional plasmid information in Table S3.

| **Optical dimerizer** | **Binding partner** | **Description** |
| --- | --- | --- |
| eMagA | eMagB, eMagBF, or eMagBM | Enhanced magnet dimerizer<sup>24</sup> |
| eMagAF | eMagB, eMagBF, or eMagBM | Enhanced magnet dimerizer with faster response kinetics<sup>24</sup> |
| eMagB | eMagA or eMagAF | Enhanced magnet dimerizer<sup>24</sup> |
| eMagBF | eMagA or eMagAF | Enhanced magnet dimerizer with faster response kinetics<sup>24</sup> |
| eMagBM | eMagA or eMagAF | Enhanced magnet dimerizer with slower response kinetics<sup>7</sup> |
| CRY2FL | CIB1 | Full length CRY2<sup>25</sup> |
| CRY2PHR | CIB1 | CRY2 truncation (residues 1-498 of CRY2)<sup>25</sup> |
| CRY2(535) | CIB1 | CRY2 truncation (residues 1-535 of CRY2)<sup>22</sup> |
| CRY2PHR (L348F) | CIB1 | Long-reversion mutant of CRY2PHR<sup>22</sup> |
| CRY2PHR<br>(W349R) | CIB1 | Short-reversion mutant of CRY2PHR<sup>22</sup> |
| EL222 | N/A (homodimerizer) | Homodimerizer with fast activation and reversion response kinetics<sup>26</sup> |
| EL222<br>(A79Q) | N/A (homodimerizer) | Medium-reversion mutant of EL222<sup>27</sup> |
| EL222<br>(AQTrip) | N/A (homodimerizer) | Long-reversion mutant of EL222<sup>27</sup> |

## Results and Discussion

### Characterization of Optogenetic Transcription Factors Using Lustro

Lustro^7^ was used to characterize the expression profiles of a set of blue light-sensitive split transcription factors (see Table 1) in response to different light induction programs. These optogenetic systems drive expression of a fluorescent protein, mScarlet-I^21^, allowing measurement of gene induction by proxy measurement of fluorescence level. Square-wave light pulses were used to induce optogenetic TFs, with varying the light pulse intensity, period, and duty cycle, where the period is the amount of time between light pulses and the duty cycle is the percentage of time the light is on during the period.. The response of each optogenetic system to this range of light induction programs is dependent on the response kinetics (activation and reversion time) of the light-sensitive proteins as well as their native light sensitivity. Relative induction level of gene expression by a TF is determined for each light condition by comparing fluorescence measurements under those conditions to the fluorescence of the constant light and constant dark control conditions for that same TF. This allows comparison of relative induction to be made between optogenetic systems, even when the magnitude of response of one system differs from another. While in-depth characterizations have been performed for a subset of optogenetic tools^22,23^, this sweep directly compares response kinetics and sensitivity of a range of optogenetic systems side-by-side in the same biological context. The maximum light pulse period within light induction programs used for screening was limited to 4 hours (with data being compared at 10 hours into induction), as longer periods will be less relevant for many applications.

### Sequential Activation of Optogenetic Systems

Data from the initial scan (Figure 1) were used to identify candidates for sequential activation, where the first light program preferentially activates one optogenetic system of a pair, and the second light program activates both optogenetic systems. Sequential activation could be useful for bioproduction processes where different stages of fermentation are desired to optimize yield^29^. CRY2(L348F)/CIB1^22,25,28^ and eMagAF/eMagBF^23,24,30^ were identified as a candidate TF pair. CRY2(L348F)/CIB1 is very sensitive to light intensity and reaches a high level of activation at low light doses. eMagAF/eMagBF is less sensitive to light intensity, requiring a higher dose of light to reach maximal activation. In order to demonstrate that sequential control of blue light systems in the same strain is possible, the TF pair was cloned into the same strain with CRY2(L348F)/CIB1 driving expression of mScarlet-I and eMagAF/eMagBF driving expression of a second, orthogonal reporter, miRFP680^31^ (and using an orthogonal DNA-binding domain, LexA^22^). Each strain was characterized in response to a range of light intensities using Lustro (see Figure 2).

**Figure 1.**
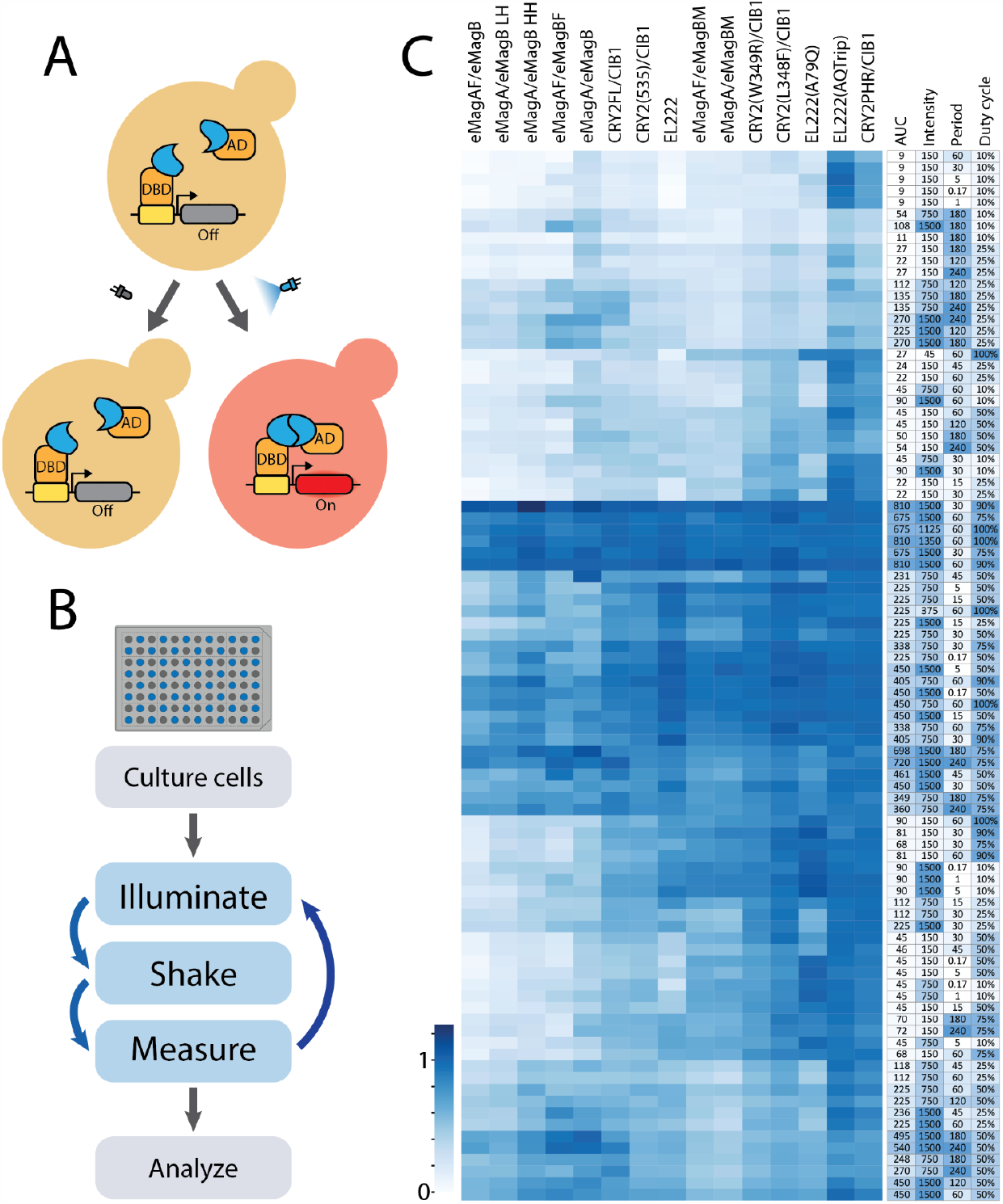
(A) Diagram showing activation of an optogenetic split TF. Blue light causes the split TF to dimerize, localizing the activation domain (AD) to the DNA-binding domain (DBD). This induces expression of the gene of interest (mScarlet-I), causing red fluorescence to increase in the cell. (B) Lustro workflow. Using laboratory automation, cells are cultured in a 96-well plate and subjected to successive rounds of illumination, shaking, and measuring, every 30 minutes. Fluorescence values are measured and analyzed. (C) Lustro was used to characterize the responses of several different optogenetic split TFs to varying light pulse intensity (μW/cm^2^), period (min), and duty cycle (%). AUC (area under the curve) is in μW·hr/cm^2^. Data shown are relative fluorescence levels (where the constant dark value is set to 0 and the constant light value is set to 1) collected 10 hours into light induction for the given TFs. LH (Low-High) and HH (High-High) designate relative expression levels of the two components of a split TF^7^ (yMM1760 and yMM1761; see Table S1), used here to demonstrate that changes in relative expression levels affect the response kinetics of two-component split TFs.

**Figure 2.**
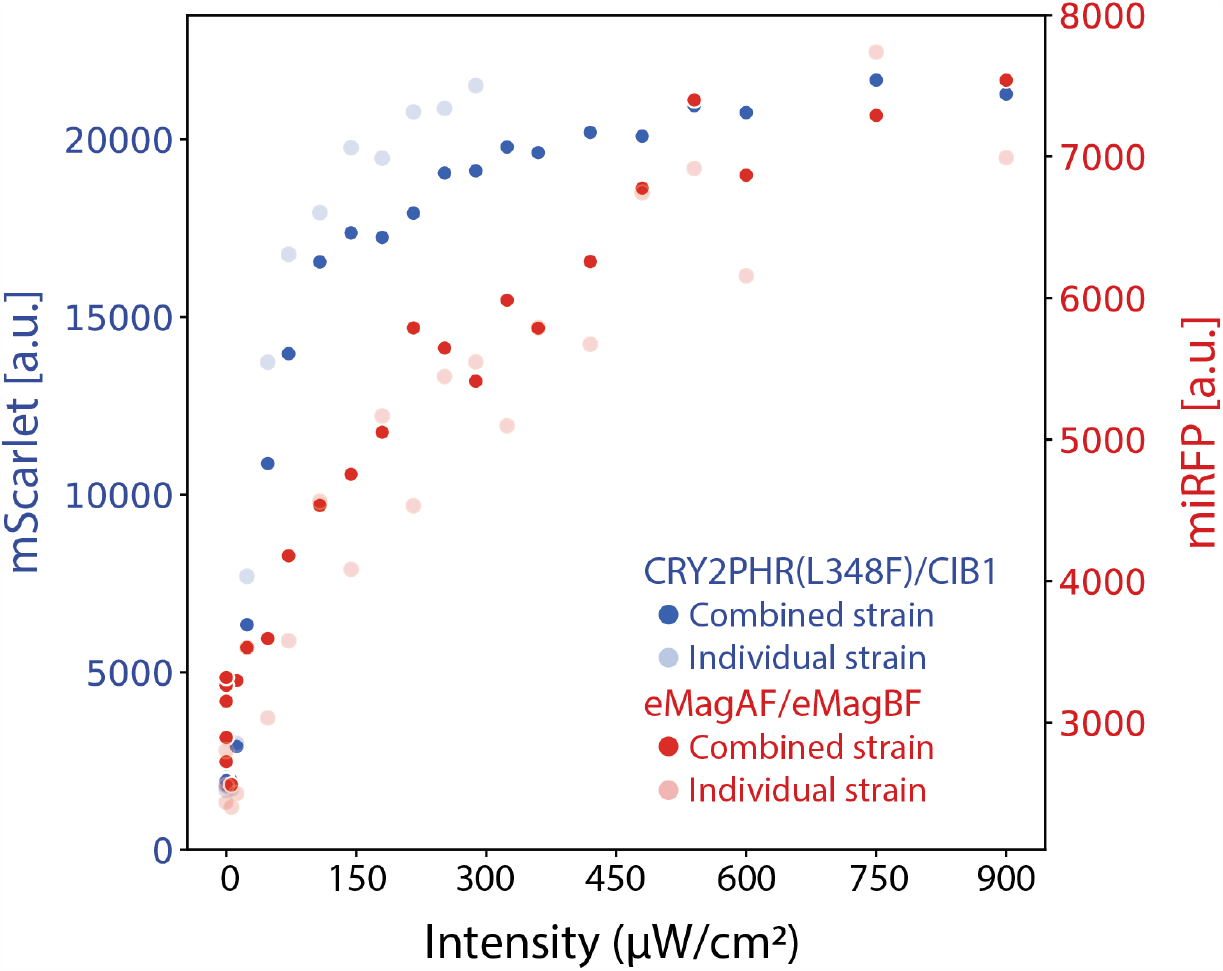
Sequential activation of optogenetic systems by a range of light intensities tested with Lustro. Two optogenetic systems are compared, CRY2PHR(L348F)/CIB1 and eMagAF/eMagBF, both engineered into the same strain (yMM1826; darker dots) and each in an individual strain (yMM1825 and yMM1781; lighter dots). The CRY2PHR(L348F)/CIB1 split TF drives expression of mScarlet (blue dots) and the eMagAF/eMagBF split TF drives expression of an orthogonal fluorescent reporter, miRFP680 (red dots). Values shown were measured after 10 hours of constant light induction. The CRY2PHR(L348F)/CIB1 system activates at lower light intensities than the eMagAF/eMagBF system.

### Multiplexed Control for Switching Between Optogenetic TFs

We next identified candidate pairs of optogenetic systems for dynamic multiplexed control over switching states^18^ (Figure 3). We took advantage of the characterization of different response kinetics and sensitivity in response to different light pulses from the initial screen performed (see Figure 1). We empirically compared relative induction between optogenetic systems and conditions in a pairwise manner to find where switching occurs. That is, where one system is activated more than another until the light induction program is changed, then the second system is activated more than the first. The 16 candidate pairs were further validated in technical quadruplicate (with a subset shown in Figure 3B). Additional switching pairs are found in Figure S2. Other useful behaviors, such as one light condition inducing both optogenetic systems to similar relative fluorescence or one optogenetic system that stays at similar relative induction between two light conditions while the other system switches, were also discovered (Figure S2). These optogenetic split TFs were characterized in separate strains, as any split TF pair combination that uses the same binding partners can freely interact and change the optogenetic activation profiles.

**Figure 3.**
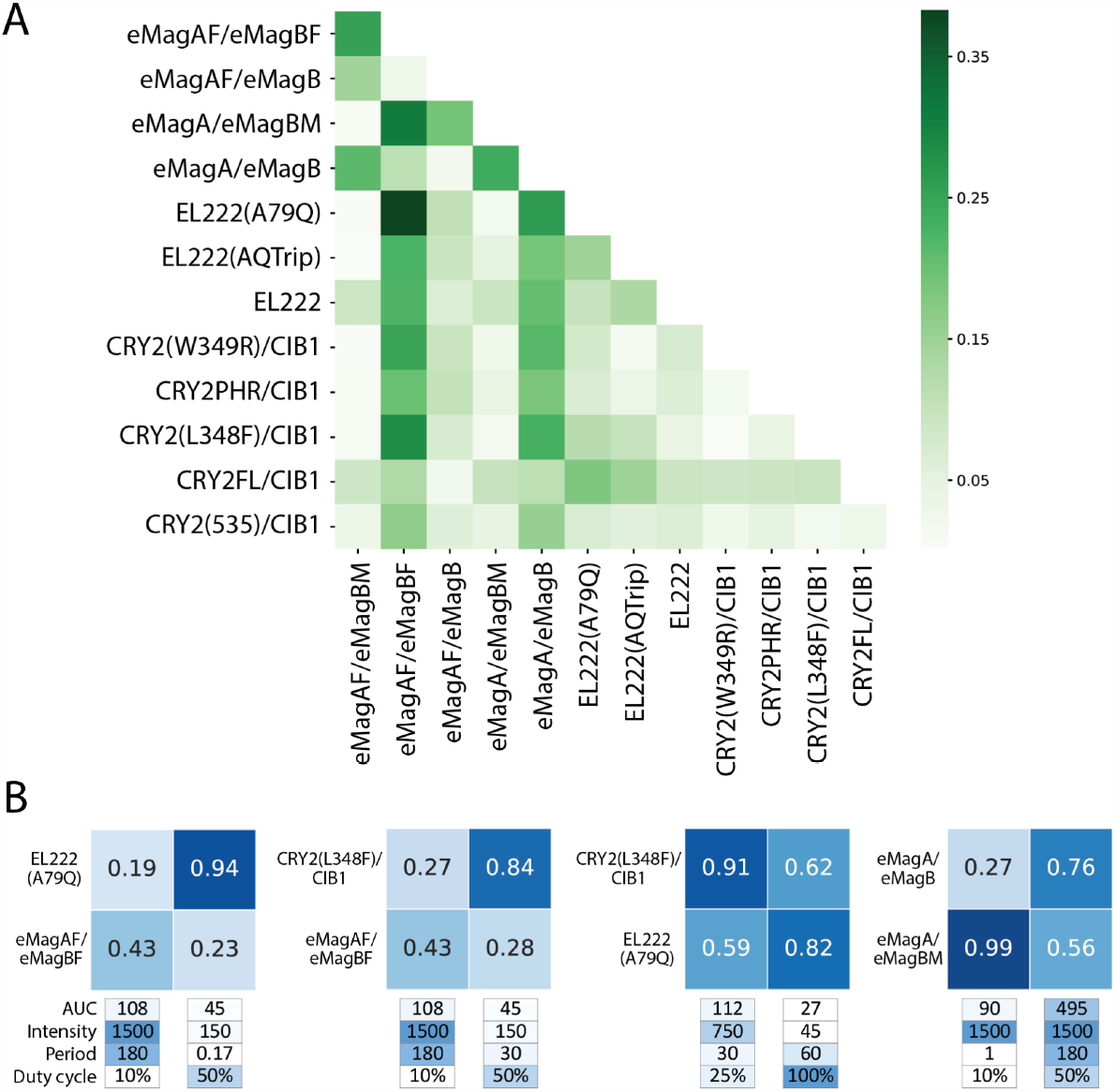
(A) Multiplexing potential for given optogenetic system pairs. First, pairwise differences between relative induction for all light conditions tested (Figure 1) for each pair of systems are calculated. The pair of light conditions that yields the highest product of differences for each pair of optogenetic systems is then calculated and plotted as a heat map (higher values of the product of differences are represented by darker blue squares). (B) Validation of pairs of optogenetic systems that switch relative induction between two different light induction conditions. Intensity is measured in μW/cm^2^, period in min, duty cycle by %, and AUC (area under the curve) is in μW·hr/cm^2^. Data shown are averaged quadruplicates of relative fluorescence, recorded at 10 hours into induction. Additional examples are presented in Figure S2.

### Response dynamics are insensitive to activation domain strength

In this study, we aimed to optimize the differences in relative response between various optogenetic systems. While controlling relative response is crucial for developing precise control strategies, we recognize that real-world applications require control over the absolute magnitude of the response. Aiming to modify the magnitude of the gene expression response of our optogenetic split TFs, we explored swapping out the activation domain. The Gal4AD in the eMagA/eMagB split TF was replaced with VP16AD, p65AD, or Msn2AD. These ADs were selected because they have been used successfully in other synthetic split TFs and have been shown to have different strengths^20,26,28,32^. Each version was characterized using Lustro, and they were found to exhibit different magnitudes of gene expression, but similar relative responses compared to each other (Figure 4). This indicates the magnitude of response changed, while the response kinetics remained similar. This experiment demonstrated the ability to adjust the magnitude of a specific optogenetic TF’s response without changing the response kinetics, thus maintaining differential responses to specific light programs.

**Figure 4.**
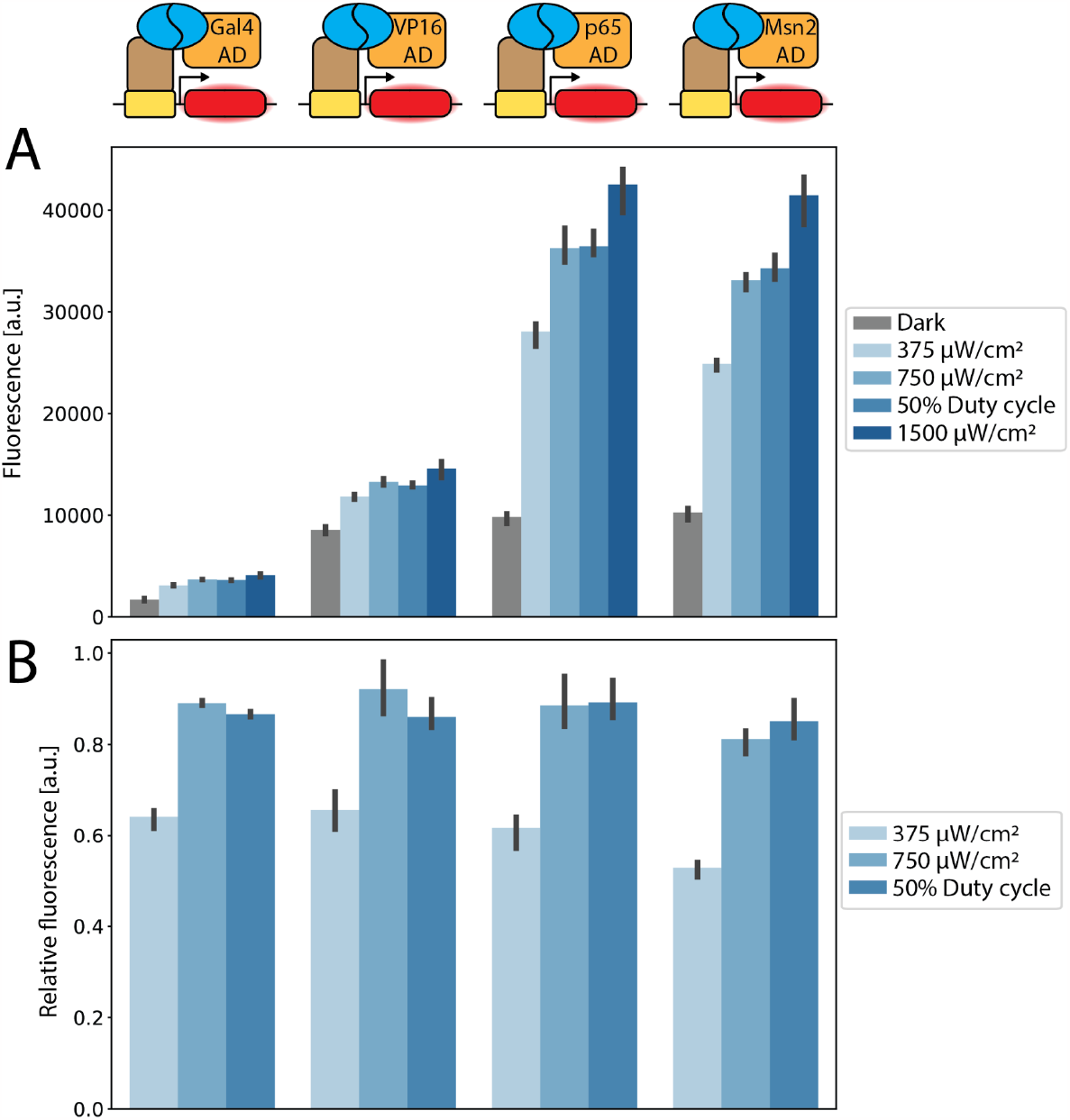
Activation domain swapping. (A) eMagA/eMagB split TF systems utilizing different activation domains are tested under a range of light conditions. Fluorescence values are shown after 10 hours of induction, with each condition performed in triplicate. The 50% duty cycle condition has an intensity of 1500 μW/cm^2^ and a period of 2 s. (B) Comparison of relative induction level (scaling each dark condition to 0 and each light condition to 1) for intermediate light induction conditions of each activation domain. An ANOVA test did not find a significant difference between relative induction levels between TFs (p > 0.5).

### Predicting System Behavior Using Machine Learning

We next sought to apply the high-throughput data collected by Lustro to generate a predictive model that would allow for selection of bespoke objective functions for various biological applications. We used a feedforward neural network (NN) to predict the relative induction of each split TF given the duty cycle, intensity, and period of the light condition. To train the neural network, we used a Bayesian inference approach to determine an approximate Gaussian distribution for the parameter posterior^33^. To evaluate model prediction performance of relative induction, we used 20-fold cross validation. This process involves dividing the data into 20 subsets, training on 19 of the subsets, and evaluating prediction performance on the held-out set. The process is repeated 20 times so that each subset is subjected to held-out testing. Prediction performance (Pearson correlation) is computed by comparing the measured relative induction to the predicted relative induction for every condition in the data set (Fig 5). The NN predicted relative induction with a Pearson correlation that ranged from 0.75 to 0.98, demonstrating that this data-driven approach provides accurate predictions of system behavior. Construction of this powerful machine learning model was enabled by the high-throughput data collection capabilities of Lustro.

**Figure 5.**
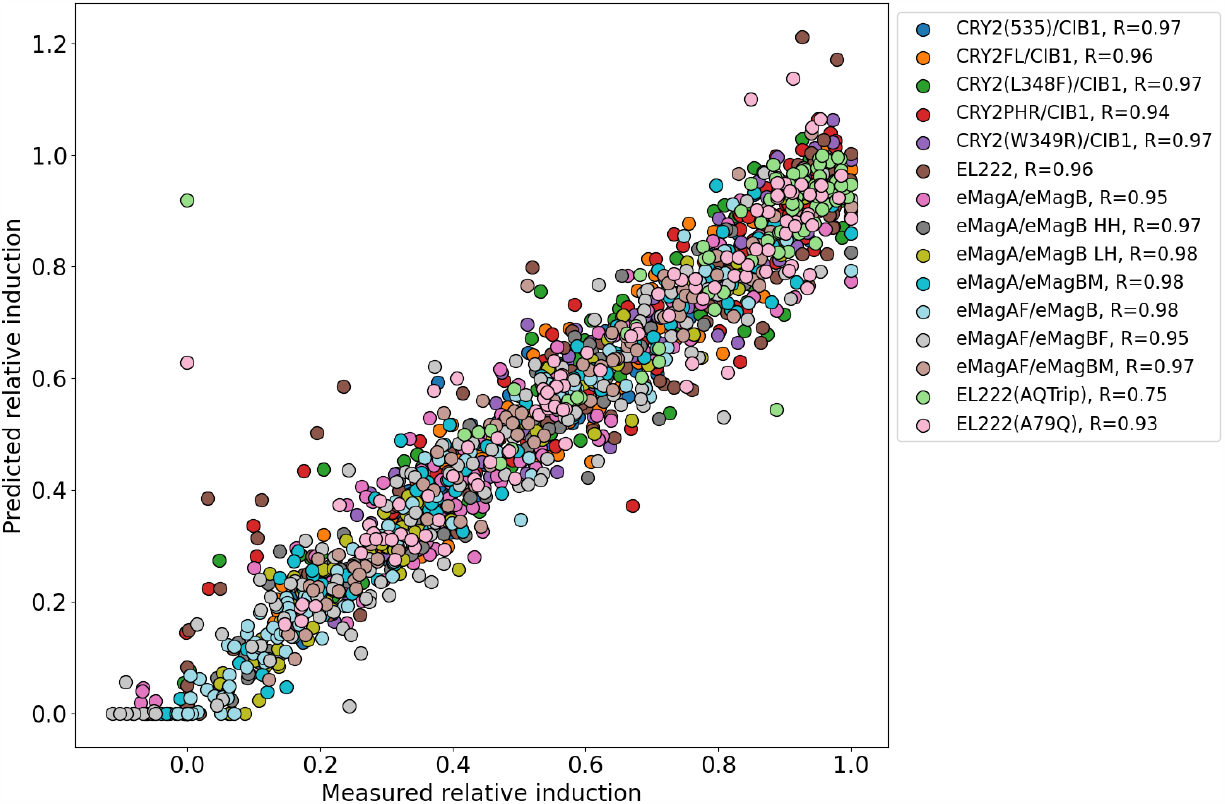
Prediction performance of relative induction using the machine learning model.

**Figure 6.**
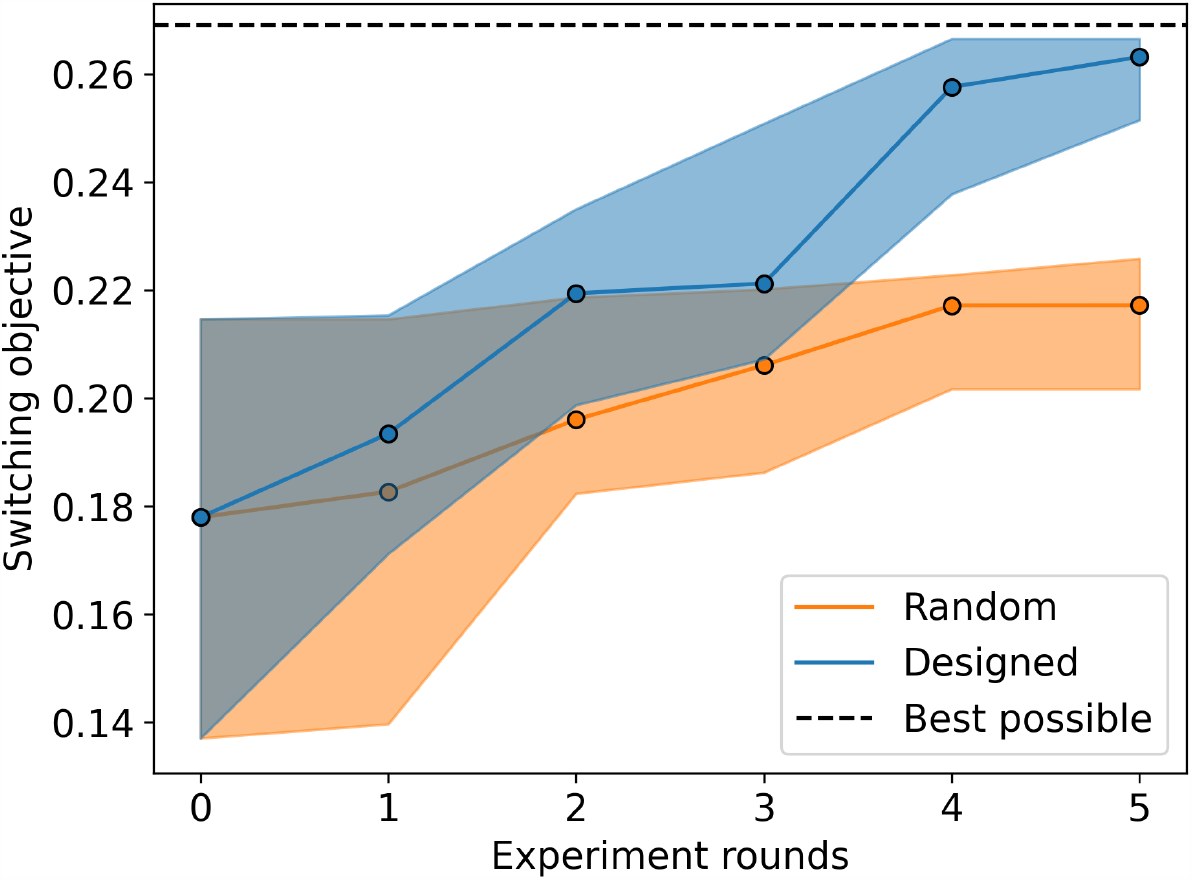
Validation of batch experimental design algorithm using the Bayesian optimization framework. Using simulated experimental data, a neural network was initialized with data from 10 randomly selected light conditions. Using the trained model, a Thompson-sampling Bayesian optimization algorithm was used to select new pairs of light conditions in subsequent experiment rounds. Compared to random selection, the model-guided experimental design algorithm more efficiently identifies conditions with improved switching in induction levels. Solid lines indicate the median performance taken over 10 trials in which the initial set of light conditions was randomly selected and shaded regions represent the interquartile range.

### Bayesian Optimization for Maximizing Switching

Once trained, the NN can be used to guide the design of experiments to select pairs of light conditions that maximize predicted switching in induction levels between two optogenetic systems. We therefore define an experimental condition as a pair of light conditions applied to a pair of split TFs. The batch data-collection capabilities of the Lustro platform enable the use of a Bayesian optimization algorithm called Thompson sampling^34^ that selects experimental conditions predicted to maximize the difference in induction between each TF pair. We use an approximate Bayesian inference approach to infer the posterior parameter distribution of the NN^33^. Once equipped with a posterior parameter distribution, Thompson sampling involves sampling parameter values from the posterior, and using the resulting model to identify the condition that maximizes the objective. The process of randomly sampling from the posterior and selecting an experimental condition that optimizes the design objective can be repeated in order to design a batch of experimental conditions. To demonstrate the potential utility of this approach, we defined a design space of pairs of light conditions scanning a range of light intensities, duty cycles, and periods. We then used a NN trained on all available experimental data to predict relative induction of all TFs for all conditions in the design space. We used these model predictions as a ‘ground truth’ dataset relating light conditions to TF induction. We then randomly selected a batch of 10 light conditions as a preliminary dataset and used this preliminary dataset to train a new NN. Using the trained model, 10 new pairs of light conditions were selected using the Thompson sampling approach to optimize an objective function defined as the negative of the product of the difference in induction levels between all pairs of TFs. The set of selected light conditions and corresponding induction levels of each TF were then queried from the ground truth dataset and appended to the training data, which was then used to update the model. The process of selecting a new set of 10 pairs of light conditions was continued over 5 rounds (each round containing a batch of experiments). The overall process was repeated over 10 trials to assess the variation in the ability of the model to optimize the system. We found that when compared to random selection of light programs, the Bayesian optimization framework quickly identified combinations of light conditions that approach the maximum possible switching in relative induction levels. These results illustrate how Lustro and machine learning can be combined to quickly identify optogenetic systems and light induction programs that enable desirable switching properties.

## Conclusion

Using a high-throughput measurement platform Lustro in combination with machine learning we described a strategy for dynamic multiplexed control of optogenetic systems that takes advantage of the native differences in response kinetics between different protein photoswitches. We used this approach to identify conditions for sequential activation and “switching” between optogenetic systems. Previous approaches integrate circuits to tune the response of an optogenetic system or generate different types of response behavior, such as OptoINVRT^35^ and OptoAMP^36^. Such circuits could be combined with this multiplexed control strategy to enable more types of optogenetic control and finer control of biological behavior. We also demonstrated the ability to tune the magnitude of response of an optogenetic system without affecting light sensitivity by changing the activation domain of the light-sensitive transcription factor.

Leveraging the predictive capabilities of a NN model, we harnessed data-driven insights to forecast the response of optogenetic systems to specific light conditions. In a simulated example, we showed that the proposed Bayesian optimization approach could rapidly identify candidate sets of TF pairs and light conditions that optimize switching in relative induction. In parallel, the implementation of workflow validation with batching represents progress towards more efficient experimental design. Reducing unnecessary iterations streamlines the process of designing and executing future experiments.

While we implemented this strategy for multiplexing light-sensitive split TFs, we propose that this method can be extended to other types of optogenetic systems. For example, this strategy could be applied to characterize and optimize other optogenetic split protein systems, such as split Cas13 systems for regulating RNA in mammalian cells^37^. The strategy could also be applied to multiplex control over optogenetic systems that regulate protein localization or oligomerization. The synergistic integration of high-throughput data collection from Lustro, NN predictive modeling, and workflow validation techniques offers a potent toolkit for advancing the frontiers of biological control. These multiplexed control strategies could be used to control bioproduction processes^35,38^, design engineered living materials^39^, regulate microbial consortia^40,41^, or interrogate complex cellular gene expression networks. Combining high-throughput characterization with machine learning to predict and optimize the behavior of optogenetic systems will rapidly accelerate the design, build, test cycle.

## Methods

### Strain Construction and Culture Conditions

Strains used in this study were constructed using standard molecular biology techniques, specifically a modular Type IIS Golden Gate assembly toolkit as previously described^4,7,42^. The details of constructs used in this work can be found in Table S3 and Table S5. Part plasmids (Level 0) were created through BsmBI Golden Gate assembly of PCR-amplified products (refer to Table S2 for primer details) or gBlocks (see Table S4) into the yTK entry vector (yTK001). Following this, part plasmids were further combined to form cassette plasmids (Level 1) using BsaI Golden Gate assembly. These cassette plasmids were then integrated into multigene plasmids (Level 2) through BsmBI Golden Gate assembly.

Single-construct strains were generated by introducing multigene plasmids, linearized with NotI-HF, into the genome of *Saccharomyces cerevisiae* strain BY4741 with the genotype MATα HIS3D1 LEU2D0 LYS2D0 URA3D0 GAL80::KANMX GAL4::spHIS5. The transformations followed an established LiAc/SS carrier DNA/PEG protocol^43^. Construct integration occurred at the URA3 or LEU2 sites, and transformants were selected using SC-Ura or SC-Leu dropout media, respectively. Transformants were further screened using previously established methods^7^.

Overnight yeast cultures were inoculated from colonies on YPD agar plates into 3 mL of liquid SC media overnight at 30 °C with agitation. Post-incubation, the overnight cultures were diluted to an optical density of 700 nm (OD700, to avoid bias from the red fluorescent marker^44^, mScarlet-I) of 0.1 in SC media. Subsequently, 200 μL of each culture was dispensed into individual wells of a 96-well glass-bottom plate with black walls (Cat. #P96-1.5H-N).

### Lustro

Automated optogenetic experiments were conducted as previously described^7,19^ using a Tecan Fluent Automation Workstation equipped with a Robotic Gripper Arm (RGA) and integrated with an optoPlate^45^, a BioShake 3000-T elm heater shaker designed for well plates, and a Tecan Spark plate reader. The optoPlate was assembled and calibrated in accordance with previously established procedures. Programming of the optoPlate was achieved using scripts available at https://github.com/mccleanlab/Optoplate-96. Throughout the experiments, the Fluent workstation was shielded from ambient light by a blackout curtain. Cellvis 96-well glass bottom plates with #1.5 cover glass (Cat. #P96-1.5H-N) were used for all experiments.

Each 96-well plate, containing cultures diluted to an optical density (OD700) of 0.1, underwent a 5-hour incubation in the dark at 30 °C with continuous shaking. Light induction commenced after this incubation period. For each light induction cycle, the plate was first positioned on the optoPlate for 26.5 minutes at 21 °C. It was then transferred to the plate shaker, where it underwent agitation at 1000 rpm with a 2 mm orbital movement for 1 minute to resuspend cells. Following this, the plate was moved to the Tecan Spark plate reader for optical density (OD700) and fluorescence measurements (without the lid). Subsequently, the plate was returned to the optoPlate, and this cycle was repeated throughout the experiment. For mScarlet-I^21^, fluorescence measurements were recorded with excitation at 563 nm and emission at 606 nm, with an optical gain of 130. For miRFP680^31^, fluorescence measurements were recorded with excitation at 652 nm and emission at 697 nm, with an optical gain of 230. The Z-value (vertical distance) was set at 28410 for all fluorescence measurements. Relative induction level for a given light induction program is calculated using the fluorescence measurements from that condition (cond), the constant light induction condition (light), and constant dark condition (dark). Relative induction = (cond - dark) / (light - dark).

### Machine Learning Model

The NN model utilized in this study is designed to predict the relative induction of each TF as a function of a particular light condition. We used a feedforward neural network architecture with a single hidden layer,

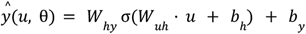

where *u* is a vector defining the light intensity, duty cycle, and the period of the light input and *y* is a vector of predicted induction levels of each TF. The parameters of the model include the weights and biases, θ = {*W*_*uh*_, *W*_*hy*_, *b*_*h*_, *b*_*y*_}.

### Bayesian Inference and Uncertainty Quantification

We used a Bayesian framework to infer a Gaussian approximation of the NN posterior parameter distribution and an Expectation-Maximization (EM) algorithm to optimize model hyperparameters, with methods adapted from Thompson et al. 2023. Model hyperparameters include the precision (inverse variance) of the parameter prior and the precision in measurement noise for each TF. The parameter prior is assumed to be a zero mean Gaussian with a precision parameter, α. We assume that error associated with measuring induction levels of *m* different TFs is a zero mean Gaussian random variable with precision β for TF *j*. Given a set of measurements of the induction levels for each TF in response to *n* different light conditions, *D* = {*y*(*u*_1_), …, *y*(*u*_*n*_)}, we define the likelihood of the data as 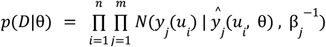. Maximizing the posterior parameter distribution with respect to model parameters is equivalent to maximizing the log of the product of the likelihood and the prior, which gives the *maximum a posteriori* (MAP) estimate, 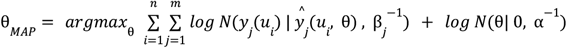. The posterior parameter distribution is approximated as a Gaussian centered at θ_*MAP*_ with a covariance matrix given by the inverse of the matrix of second derivatives of the negative log posterior, which we approximate using the outer product, 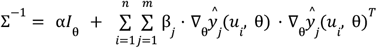 where *I*_θ_ is the identity matrix with dimension equal to the number of model parameters. The model hyperparameters α and β are optimized using the EM algorithm, which involves maximizing the expectation of the log of the joint probability of the data and the parameter distribution with respect to α and β, where the expectation is taken with respect to the parameter posterior distribution. Using the updated hyperparameters, inference of the posterior parameter distribution is repeated until convergence of the marginal likelihood.

### Experimental Design using Bayesian Optimization

We used a Bayesian optimization algorithm called Thompson sampling to design a batch of experimental conditions predicted to maximize the difference in induction levels between pairs of TFs in separate light conditions. To do so, we define the objective function as the negative of the minimum product in the difference between predicted induction for each TF,

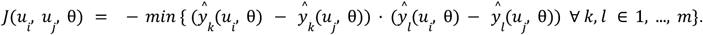

We define the experimental design space as all possible pairs of light conditions, *Q* = { (*u*_*i*_, *u*_*j*_) ∀ *i* ≠ *j*}. The Thompson sampling algorithm involves sampling parameter values from the posterior, θ^*^ ∼ *N*(θ_*MAP*_, Σ), and then determining the experimental condition that maximizes the objective, 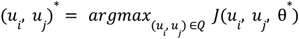. This process is repeated as many times as necessary to generate the desired number of experimental conditions to be tested in the next experiment.

## Materials Availability

Key plasmids have been deposited on Addgene. For all other reagent requests, please contact the corresponding author.

## Supporting Information

Additional data and schematics for the experiments described in the text, strains, plasmids, oligos, gene blocks, and optogenetic constructs used in this work

## Acknowledgments

This work was supported by National Institutes of Health grant R35GM128873 and National Science Foundation grant 2045493 (awarded to M.N.M.) and National Science Foundation CBET 2315963 (awarded to V.M.Z.) Megan Nicole McClean, PhD, holds a Career Award at the Scientific Interface from the Burroughs Wellcome Fund. Z.P.H. was supported by an NHGRI training grant to the Genomic Sciences Training Program 5T32HG002760. We thank Amit Nimunkar and Edvard Grødem for building and modifying the optoPlate. We acknowledge fruitful discussions with McClean lab members, Neydis Moreno for providing feedback on the manuscript, and Stephanie Geller for providing pMM1134 and pMM1137.

## Author Contributions

Z.P.H. and M.N.M conceived of the study. J.C.T., D.L.C., and V.M.Z. conceived of the modeling approach. Z.P.H. designed optogenetic parts, performed experiments, and analyzed data. J.C.T. designed the neural network and generated the predictive model. M.N.M. and V.M.Z. provided funding. Z.P.H. and J.C.T. wrote the original draft of the manuscript, and Z.P.H., J.C.T., D.L.C., V.M.Z. and M.N.M wrote, reviewed, and edited the final manuscript.

