## Supplemental for "Dynamic Multiplexed Control and Modeling of Optogenetic Systems Using the High-Throughput Optogenetic Platform, Lustro"

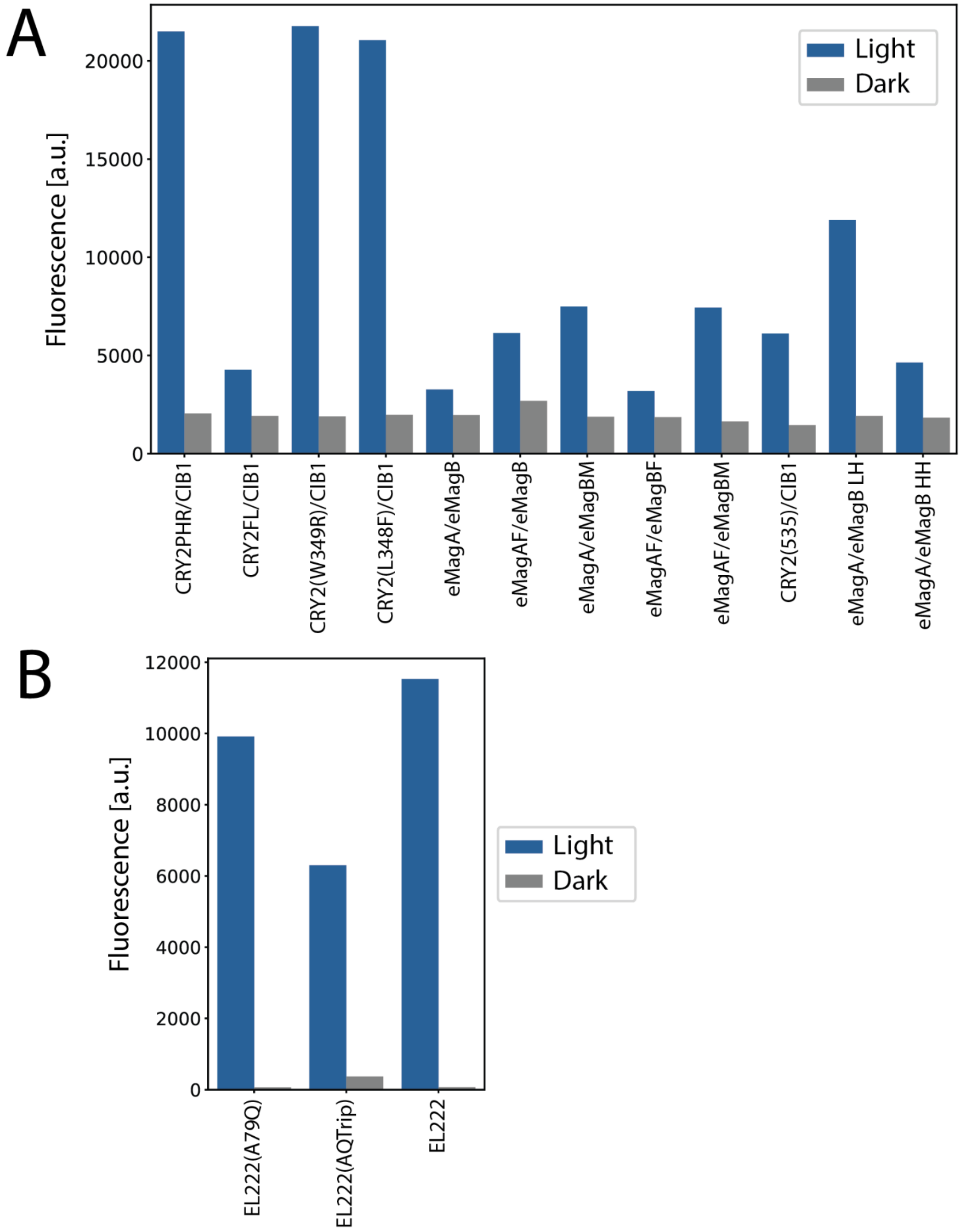


**Figure S1.** Magnitude of fluorescence response for optogenetically induced strains measured in Figure 1. The fluorescence values shown here for the light and dark conditions are set to 1 and 0, respectively, to calculate relative activation levels. Fluorescence values shown are recorded at 10 hours of induction. (A) Strains measured with 130 optical gain. (B) Strains measured with 80 optical gain.

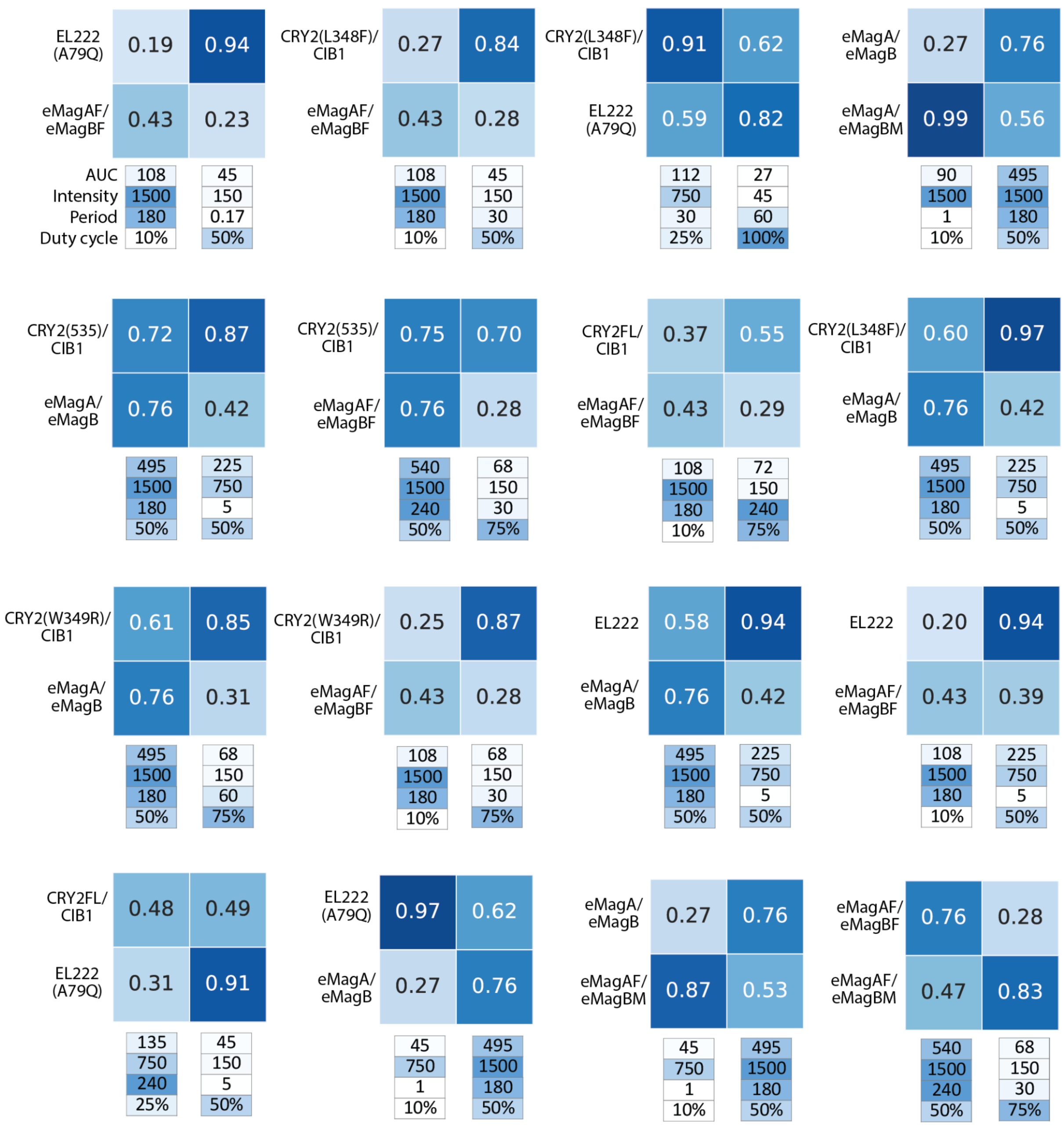


**Figure S2.** Extension of Figure 3. Validation of pairs of optogenetic systems that switch relative activation amounts between two different light induction conditions. While most of the pairs exhibit switching behavior, several other interesting types of behavior are observed, such as one light condition holding both strains at similar relative activation (CRY2(535)/CIB1 and eMagA/eMagB) or one strain staying at similar relative activation between two light conditions while the other strain switches (CRY2FL/CIB1 and EL222(A79Q)). Intensity is measured in μW/cm$^2$, period in min, duty cycle by %, and AUC (area under the curve) is in μW·hr/cm$^2$. Data shown are averaged quadruplicates of relative fluorescence, recorded at 10 hours into induction.

Table S1: Yeast strains used in this study

| ID | Alias | Genotype | Description | Source |
|---|---|---|---|---|
| yMM1731 | pRPL18B-Gal4DBD-CRY2PHR, pRPL18B-Gal4AD-CIB1, pGAL1-mScarlet-I | BY4741 Matα ura3Δ0::5' Ura3 homology, pRPL18B-Gal4DBD-CRY2PHR-tENO1, pRPL18B-Gal4AD-CIB1-tENO1, pGAL1-mScarlet-I-tENO1, Ura3, Ura 3' homology his3D1 leu2D0 lys2D0 gal80::KANMX gal4::spHIS5 | pRPL18B-CRY2PHR, pRPL18B-CIB1 | Harmer et al. 2023 ACS Syn Bio[1] |
| yMM1733 | pRPL18B-Gal4DBD-CRY2, pRPL18B-Gal4AD-CIB1, pGAL1-mScarlet-I | BY4741 Matα ura3Δ0::5' Ura3 homology, pRPL18B-Gal4DBD-CRY2-tENO1, pRPL18B-Gal4AD-CIB1-tENO1, pGAL1-mScarlet-I-tENO1, Ura3, Ura 3' homology his3D1 leu2D0 lys2D0 gal80::KANMX gal4::spHIS5 | pRPL18B-CRY2FL, pRPL18B-CIB1 | Harmer et al. 2023 ACS Syn Bio[1] |
| yMM1734 | pRPL18B-Gal4DBD-eMagA, pRPL18B-eMagB-Gal4AD, pGAL1-mScarlet-I | BY4741 Matα ura3Δ0::5' Ura3 homology, pRPL18B-Gal4DBD-eMagA-tENO1, pRPL18B-eMagB-Gal4AD-tENO1, pGAL1-mScarlet-I-tENO1, Ura3, Ura 3' homology his3D1 leu2D0 lys2D0 gal80::KANMX gal4::spHIS5 | pRPL18B-eMagA, pRPL18B-eMagB | Harmer et al. 2023 ACS Syn Bio[1] |
| yMM1760 | pRPL18B-Gal4DBD-eMagA, pTEF1-eMagB-Gal4AD, pGAL1-mScarlet-I (LH) | BY4741 Matα ura3Δ0::5' Ura3 homology, pRPL18B-Gal4DBD-eMagA-tENO1, pTEF1-eMagB-Gal4AD-tENO1, pGAL1-mScarlet-I-tENO1, Ura3, Ura 3' homology KanR-ColE1 his3D1 leu2D0 lys2D0 gal80::KANMX gal4::spHIS5 | pRPL18B-eMagA, pTEF1-eMagB | Harmer et al. 2023 ACS Syn Bio[1] |
| yMM1761 | pTEF1-Gal4DBD-eMagA, pTEF1-eMagB-Gal4AD, pGAL1-mScarlet-I (HH) | BY4741 Matα ura3Δ0::5' Ura3 homology, pTEF1-Gal4DBD-eMagA-tENO1, pTEF1-eMagB-Gal4AD-tENO1, pGAL1-mScarlet-I-tENO1, Ura3, Ura 3' homology his3D1 leu2D0 lys2D0 gal80::KANMX gal4::spHIS5 | pTEF1-eMagA, pTEF1-eMagB | Harmer et al. 2023 ACS Syn Bio[1] |
| yMM1762 | pCCW12-NLS-VP16-EL222-tENO1, spacer, 5xBS | BY4741 Matα ura3Δ0::5' Ura3 homology, pCCW12-NLS-VP16-EL222-tENO1, spacer, 5xBS | pCCW12-NLS-VP16-EL222 | This work |

| | | | | |
|---|---|---|---|---|
| | CYC180pr-mScarlet-I | CYC180pr-mScarlet-I-tENO1, Ura3, Ura 3' homology his3D1 leu2D0 lys2D0 gal80::KANMX gal4::spHIS5 | | |
| yMM1763 | pRPL18B-Gal4DBD-CRY2(535), pRPL18B-Gal4AD-CIB1, pGAL1-mScarlet-I | BY4741 Matα ura3Δ0::5' Ura3 homology, pRPL18B-Gal4DBD-CRY2(535)-tENO1, pRPL18B-Gal4AD-CIB1-tENO1, pGAL1-mScarlet-I-tENO1, Ura3, Ura 3' homology his3D1 leu2D0 lys2D0 gal80::KANMX gal4::spHIS5 | pRPL18B-CRY2(535), pRPL18B-CIB1 | Harmer et al. 2023 ACS Syn Bio[1] |
| yMM1764 | pRPL18B-Gal4DBD-eMagAF, pRPL18B-eMagBF-Gal4AD, pGAL1-mScarlet-I | BY4741 Matα ura3Δ0::5' Ura3 homology, pRPL18B-Gal4DBD-eMagAF-tENO1, pRPL18B-eMagBF-Gal4AD-tENO1, pGAL1-mScarlet-I-tENO1, Ura3, Ura 3' homology his3D1 leu2D0 lys2D0 gal80::KANMX gal4::spHIS5 | pRPL18B-eMagAF, pRPL18B-eMagBF | Harmer et al. 2023 ACS Syn Bio[1] |
| yMM1765 | pRPL18B-Gal4DBD-eMagA, pRPL18B-eMagBM-Gal4AD, pGAL1-mScarlet-I | BY4741 Matα ura3Δ0::5' Ura3 homology, pRPL18B-Gal4DBD-eMagA-tENO1, pRPL18B-eMagBM-Gal4AD-tENO1, pGAL1-mScarlet-I-tENO1, Ura3, Ura 3' homology his3D1 leu2D0 lys2D0 gal80::KANMX gal4::spHIS5 | pRPL18B-eMagA, pRPL18B-eMagBM | Harmer et al. 2023 ACS Syn Bio[1] |
| yMM1777 | pRPL18B-Gal4BD-eMagAF-tENO1, pRPL18B-eMagBM-Gal4AD-tENO1, pGAL1-mScarlet-I-tENO1, Ura3 | BY4741 Matα ura3Δ0::5' Ura3 homology, pRPL18B-Gal4BD-eMagAF-tENO1, pRPL18B-eMagBM-Gal4AD-tENO1, pGAL1-mScarlet-I-tENO1, Ura3, Ura 3' homology his3D1 leu2D0 lys2D0 gal80::KANMX gal4::spHIS5 | pRPL18B-eMagAF, pRPL18B-eMagBM | This work |
| yMM1778 | pRPL18B-Gal4BD-eMagAF-tENO1, pRPL18B-eMagB-Gal4AD-tENO1, pGAL1-mScarlet-I-tENO1, Ura3 | BY4741 Matα ura3Δ0::5' Ura3 homology, pRPL18B-Gal4BD-eMagAF-tENO1, pRPL18B-eMagB-Gal4AD-tENO1, pGAL1-mScarlet-I-tENO1, Ura3, Ura 3' homology his3D1 leu2D0 lys2D0 gal80::KANMX gal4::spHIS5 | pRPL18B-eMagAF, pRPL18B-eMagB | This work |
| yMM1780 | pRPL18B-Gal4BD-CRY2PHR | BY4741 Matα ura3Δ0::5' Ura3 homology, | pRPL18B-CRY2PHR(W | This work |

| | | | | |
|---|---|---|---|---|
| | W349R-tENO1, pRPL18B-Gal4AD-CIB1-tENO1, pGAL1-mScarlet-I-tENO1, Ura3 | pRPL18B-Gal4BD-CRY2PHRW349R-tENO1, pRPL18B-Gal4AD-CIB1-tENO1, pGAL1-mScarlet-I-tENO1, Ura3, Ura 3' homology his3D1 leu2D0 lys2D0 gal80::KANMX gal4::spHIS5 | 349R), pRPL18B-CIB1 | |
| yMM1781 | pRPL18B-Gal4BD-CRY2PHRL348F-tENO1, pRPL18B-Gal4AD-CIB1-tENO1, pGAL1-mScarlet-I-tENO1, Ura3 | BY4741 Matα ura3Δ0::5' Ura3 homology, pRPL18B-Gal4BD-CRY2PHRL348F-tENO1, pRPL18B-Gal4AD-CIB1-tENO1, pGAL1-mScarlet-I-tENO1, Ura3, Ura 3' homology his3D1 leu2D0 lys2D0 gal80::KANMX gal4::spHIS5 | pRPL18B-CRY2PHR(L348F), pRPL18B-CIB1 | This work |
| yMM1812 | pCCW12-NLS-VP16-EL222(A79Q)-tENO1, spacer, 5xBS CYC180pr-mScarlet-I-tENO1, Ura3 | BY4741 Matα ura3Δ0::5' Ura3 homology, pCCW12-NLS-VP16-EL222(A79Q)-tENO1, spacer, 5xBS CYC180pr-mScarlet-I-tENO1, Ura3, Ura 3' homology his3D1 leu2D0 lys2D0 gal80::KANMX gal4::spHIS5 | pCCW12-NLS-VP16-EL222(A79Q) | This work |
| yMM1813 | pCCW12-NLS-VP16-EL222AQTrip-tENO1, spacer, 5xBS CYC180pr-mScarlet-I-tENO1, Ura3 | BY4741 Matα ura3Δ0::5' Ura3 homology, pCCW12-NLS-VP16-EL222AQTrip-tENO1, spacer, 5xBS CYC180pr-mScarlet-I-tENO1, Ura3, Ura 3' homology his3D1 leu2D0 lys2D0 gal80::KANMX gal4::spHIS5 | pCCW12-NLS-VP16-EL222(AQTrip) | This work |
| yMM1819 | pRPL18B-Gal4DBD-eMagA-tENO1, pRPL18B-eMagB-p65AD-tENO1, pGAL1-mScarlet-I-tENO1, Ura3 | BY4741 Matα ura3Δ0::5' Ura3 homology, pRPL18B-Gal4DBD-eMagA-tENO1, pRPL18B-eMagB-p65AD-tENO1, pGAL1-mScarlet-I-tENO1, Ura3, Ura 3' homology his3D1 leu2D0 lys2D0 gal80::KANMX gal4::spHIS5 | pRPL18B-eMagA, pRPL18B-eMagB-p65AD | This work |
| yMM1820 | pRPL18B-Gal4DBD-eMagA-tENO1, pRPL18B-eMagB-Msn2AD-tENO1, | BY4741 Matα ura3Δ0::5' Ura3 homology, pRPL18B-Gal4DBD-eMagA-tENO1, pRPL18B-eMagB-Msn2AD-tENO1, pGAL1-mScarlet-I-tENO1, Ura3, Ura 3' homology his3D1 leu2D0 lys2D0 | pRPL18B-eMagA, pRPL18B-eMagB-Msn2AD | This work |

| | | | | |
|---|---|---|---|---|
| | pGAL1-mScarlet-I-tENO1, Ura3 | gal80::KANMX gal4::spHIS5 | | |
| yMM1821 | pRPL18B-Gal4DBD-eMagA-tENO1, pRPL18B-eMagB-VP16AD-tENO1, pGAL1-mScarlet-I-tENO1, Ura3 | BY4741 Matα ura3Δ0::5' Ura3 homology, pRPL18B-Gal4DBD-eMagA-tENO1, pRPL18B-eMagB-VP16AD-tENO1, pGAL1-mScarlet-I-tENO1, Ura3, Ura 3' homology his3D1 leu2D0 lys2D0 gal80::KANMX gal4::spHIS5 | pRPL18B-eMagA, pRPL18B-eMagB-VP16AD | This work |
| yMM1825 | pRPL18B-LexA-eMagAF-tENO1, pRPL18B-eMagBF-Gal4AD-tENO1, pLexA-miRFP680-tENO1, Leu2 | BY4741 Matα leu2Δ0::5' Ura3 homology, pRPL18B-LexA-eMagAF-tENO1, pRPL18B-eMagBF-Gal4AD-tENO1, pLexA-miRFP680-tENO1, Leu2, Leu2 3' homology his3D1 ura3D0 lys2D0 gal80::KANMX gal4::spHIS5 | pRPL18B-eMagAF, pRPL18B-eMagBF, pLexA-miRFP680 | This work |
| yMM1826 | pRPL18B-Gal4BD-CRY2PHRL348F-tENO1, pRPL18B-Gal4AD-CIB1-tENO1, pGAL1-mScarlet-I-tENO1, Ura3; pRPL18B-LexA-eMagAF-tENO1, pRPL18B-eMagBF-Gal4AD-tENO1, pLexA-miRFP680-tENO1, Leu2 | BY4741 Matα ura3Δ0::5' Ura3 homology, pRPL18B-Gal4BD-CRY2PHRL348F-tENO1, pRPL18B-Gal4AD-CIB1-tENO1, pGAL1-mScarlet-I-tENO1 Ura3, Ura 3' homology; leu2D0::5' Leu2 homology, pRPL18B-LexA-eMagAF-tENO1, pRPL18B-eMagBF-Gal4AD-tENO1, pLexA-miRFP680-tENO1, Leu2, Leu2 3' homology his3D1 lys2D0 gal80::KANMX gal4::spHIS5 | pRPL18B-CRY2PHR(L348F), pRPL18B-CIB1, pGAL1-mScarlet-I; pRPL18B-eMagAF, pRPL18B-eMagBF, pLexA-miRFP680 | This work |

#### Table S2: Primers used in this study

| **ID** | **Alias** | **Sequence** | **Target** | **Purpose** |
|---|---|---|---|---|

| | | | | |
|---|---|---|---|---|
| oMM2321 | Fwd Q5 L348F CRY2PHR | AATGAGAGAGtTTTGGGCTACCGG | CRY2PHR | Q5 SDM to make CRY2PHR(L348F) |
| oMM2322 | Rev Q5 L348F CRY2PHR | CCGGCATCCACCAACGGA | CRY2PHR | Q5 SDM to make CRY2PHR(L348F) |
| oMM2323 | Fwd Q5 W349R CRY2PHR | GAGAGAGCTTaGGGCTACCGG | CRY2PHR | Q5 SDM to make CRY2PHR(W349R) |
| oMM2324 | Rev Q5 W349R CRY2PHR | ATTCCGGCATCCACCAAC | CRY2PHR | Q5 SDM to make CRY2PHR(W349R) |
| oMM2445 | 5xBS-CYC180pr insert 1 | gcatCGTCTCaTCGGTCTCAAACGGGGAGATCTTCGCTAGCCTCGAGTAtGAGACGgcat | | Golden Gate assembly of 5xBS-CYC180pr |
| oMM2446 | 5xBS-CYC180pr insert 2 | gcatCGTCTCaAGTAGGTAGCCTTTAGTCCATGCGTTATAGGTAGCCTTtGAGACGgcat | | Golden Gate assembly of 5xBS-CYC180pr |
| oMM2447 | 5xBS-CYC180pr insert 3 | gcatCGTCTCaCCTTTAGTCCATGCGTTATAGGTAGCCTTTAGTCCATGtGAGACGgcat | | Golden Gate assembly of 5xBS-CYC180pr |
| oMM2448 | 5xBS-CYC180pr insert 4 | gcatCGTCTCaCATGCGTTATAGGTAGCCTTTAGTCCATGCGTTATAGtGAGACGgcat | | Golden Gate assembly of 5xBS-CYC180pr |
| oMM2464 | 5xBS-CYC180pr insert 1 RC | atgcCGTCTCaTACTCGAGGCTAGCGAAGATCTCCCCGTTTGAGACCGAtGAGACGatgc | | Golden Gate assembly of 5xBS-CYC180pr |

| | | | | |
|---|---|---|---|---|
| oMM2465 | 5xBS-CYC180pr insert 2 RC | atgcCGTCTCaAAGGCTACCTATAACGCATGGACTAAAGGCTACCTACTtGAGACGatgc | | Golden Gate assembly of 5xBS-CYC180pr |
| oMM2466 | 5xBS-CYC180pr insert 3 RC | atgcCGTCTCaCATGGACTAAAGGCTACCTATAACGCATGGACTAAAGGtGAGACGatgc | | Golden Gate assembly of 5xBS-CYC180pr |
| oMM2467 | 5xBS-CYC180pr insert 4 RC | atgcCGTCTCaCTATAACGCATGGACTAAAGGCTACCTATAACGCATGtGAGACGatgc | | Golden Gate assembly of 5xBS-CYC180pr |
| oMM2781 | Fwd EL222 A79Q | CCGATTCCTGcaAGGTTCCGGCACc | EL222 | Q5 SDM to make EL222A79Q from pMM1176 |
| oMM2782 | Rev EL222 A79Q | CAATTGCGGCCGACGCAT | EL222 | Q5 SDM to make EL222A79Q from pMM1176 |
| oMM2796 | Fwd msn2AD 3b | gcatCGTCTCatcggtctcaTTCTggccctaaaaagaagcgtaaagtcACGGTCGACCATGATTTC | Msn2AD | Amplify NLS-msn2AD 3b insert from pMM0565 |
| oMM2798 | Fwd VP16AD 3b | gcatCGTCTCatcggtctcaTTCTGGCCCTAAAAAGAAGCGTAAAG | VP16AD | Amplify VP16 3b from pMM1432 |
| oMM2799 | Rev VP16AD 3b | atgcCGTCTCaggtctcaGGATttaCCCACCGTACTCGTCAATTC | VP16AD | Amplify VP16 3b from pMM1432 |
| oMM2807 | Rev msn2AD 3b | atgcCGTCTCaggtctcaGGATttaGTTTGTTATAACGTCGCTAAA | Msn2AD | Amplify NLS-msn2AD 3b insert from pMM0565 |

#### Table S3: Plasmids used in this study

| ID | Alias | Gene(s) or Insert | Yeast Marker | Bacterial Resistance | Toolkit Part | Source |
|---|---|---|---|---|---|---|
| pMM0452 | Entry vector (pYTK001; Addgene #65108) | ColE1-CamR-sfGFP dropout | | CamR | Entry vector | Lee et al. 2015 Acs Syn Bio[2] |
| pMM0454 | pRPL18B (pYTK017; Addgene #65124) | ColE1-CamR-pRPL18B | | CamR | Part, Type 2 | Lee et al. 2015 Acs Syn Bio[2] |
| pMM0458 | GFP dropout cassette (pYTK096) | ColE1-KanR-URA3 3' homology-URA3-sfGFP dropout-URA3 5' homology | URA3 | KanR | GFP dropout cassette | Lee et al. 2015 Acs Syn Bio[2] |
| pMM0477 | ConLS' (pYTK008; Addgene #65115) | ColE1-CamR-ConLS' | | CamR | Part, Type 1 | Lee et al. 2015 Acs Syn Bio[2] |
| pMM0478 | ConRE' (pYTK073; Addgene #65180) | ColE1-CamR-ConRE' | | CamR | Part, Type 5 | Lee et al. 2015 Acs Syn Bio[2] |
| pMM0479 | LEU2 (pYTK075; Addgene #65182) | ColE1-CamR-LEU2 | | CamR | Part, Type 6 | Lee et al. 2015 Acs Syn Bio[2] |
| pMM0480 | LEU2 3' homology (pYTK087; Addgene #65194) | ColE1-CamR-LEU2 3' homology | | CamR | Part, Type 7 | Lee et al. 2015 Acs Syn Bio[2] |
| pMM0481 | RFP (pYTK090; Addgene #65197) | ColE1-KanR-RFP | | CamR | Part, Type 8a | Lee et al. 2015 Acs Syn Bio[2] |

| | | | | | | |
|---|---|---|---|---|---|---|
| pMM0482 | LEU2 5' homology (pYTK093; Addgene #65200) | ColE1-CamR-LEU2 5' homology | | CamR | Part, Type 8b | Lee et al. 2015 Acs Syn Bio[2] |
| pMM0489 | ConLS (pYTK002; Addgene #65109) | ColE1-CamR-ConLS | | CamR | Part, Type 1 | Lee et al. 2015 Acs Syn Bio[2] |
| pMM0490 | sfGFP dropout (pYTK047; Addgene #65154) | ColE1-CamR-sfGFP dropout | | CamR | Part, Type 234 | Lee et al. 2015 Acs Syn Bio[2] |
| pMM0491 | ConR1 (pYTK067; Addgene #65174) | ColE1-CamR-ConR1 | | CamR | Part, Type 5 | Lee et al. 2015 Acs Syn Bio[2] |
| pMM0532 | ConL1 (pYTK003; Addgene #65110) | ColE1-CamR-ConL1 | | CamR | Part, Type 1 | Lee et al. 2015 Acs Syn Bio[2] |
| pMM0533 | ConL2 (pYTK004; Addgene #65111) | ColE1-CamR-ConL2 | | CamR | Part, Type 1 | Lee et al. 2015 Acs Syn Bio[2] |
| pMM0537 | ConR2 (pYTK068; Addgene #65175) | ColE1-CamR-ConR2 | | CamR | Part, Type 5 | Lee et al. 2015 Acs Syn Bio[2] |
| pMM0541 | ConRE (pYTK072; Addgene #65179) | ColE1-CamR-ConRE | | CamR | Part, Type 5 | Lee et al. 2015 Acs Syn Bio[2] |
| pMM0542 | tENO1 (pYTK051; Addgene #65158) | ColE1-CamR-tENO1 | | CamR | Part, Type 4 | Lee et al. 2015 Acs Syn Bio[2] |
| pMM0556 | sfGFP dropout (pYTK095; Addgene | ColE1-AmpR-sfGFP dropout | | AmpR | 678 | Lee et al. 2015 Acs Syn Bio[2] |

| | | | | | | |
|---|---|---|---|---|---|---|
| | #65202) | | | | | |
| pMM0559 | pCCW12 (pYTK010; Addgene #65117) | ColE1-CamR-pCCW12 | | CamR | Part, Type 2 | Lee et al. 2015 Acs Syn Bio[2] |
| pMM0565 | pTEF-wtMsn2-mCherry-LINuS | pTEF-wtMsn2-mCherry-LINuS | | AmpR | | Niopek et al. 2014 Nat Comm[3] |
| pMM0777 | Spacer | ConL1-Spacer-ConR2-Leu2-CEN/ARS-ColE1-KanR | | AmpR | Cassette | Geller et al. 2019 Cell Mol Bioeng[4] |
| pMM0918 | Gal4DBD (n-terminal) | ColE1-CamR-Gal4DBD | | CamR | Part, Type 3a | Harmer et al. 2023 ACS Syn Bio[1] |
| pMM0921 | CRY2PHR | ColE1-CamR-CRY2PHR | | CamR | Part, Type 3b | Harmer et al. 2023 ACS Syn Bio[1] |
| pMM0940 | miRFP680 | ColE1-CamR-miRFP680 | | CamR | Part, Type 3 | Matlashov et al. 2020 Nat Comm[5]; This work |
| pMM0946 | pRPL18B-Gal4AD-CIB1 cassette | ConL1-pRPL18B-Gal4AD-CIB1-tENO1-ConR2 | | AmpR | Cassette | Harmer et al. 2023 ACS Syn Bio[1] |
| pMM1088 | CRY2PHR L348F | ColE1-CamR-CRY2PHR L348F | | CamR | Part, Type 3b | This work |
| pMM1089 | CRY2PHR W349R | ColE1-CamR-CRY2PHR W349R | | CamR | Part, Type 3b | This work |
| pMM1090 | pRPL18B-Gal4DBD-CRY2PHRL348F | ConLS-pRPL18B-Gal4DBD-CRY2PHRL348F-tENO1-ConR1 | | AmpR | Cassette | This work |
| pMM1091 | pRPL18B-Gal4DBD-CR | ConLS-pRPL18B-Gal4DBD-CRY2PHRW3 | | AmpR | Cassette | This work |

| | | | | | | |
|---|---|---|---|---|---|---|
| | Y2PHRW349R | 49R-tENO1-ConR2 | | | | |
| pMM1098 | 5xBS CYC180pr | ColE1-CamR-5xBS CYC180pr | | CamR | Part, Type 2 | Benzinger et al. 2018 Nat Comm[6]; This work |
| pMM1134 | LexA | ColE1-CamR-LexA | | CamR | Part, Type 3 | This work |
| pMM1137 | pLexA | ColE1-CamR-pLexA | | CamR | Part, Type 2 | This work |
| pMM1176 | NLS-VP16-EL222 | ColE1-CamR-NLS-VP16-EL222 | | CamR | Part, Type 3 | Motta-Mena et al. 2014 Nat Chem Biol[7]; This work |
| pMM1177 | pCCW12-NLS-VP16-EL222 | ConLS-pCCW12-NLS-VP16-EL222-tENO1-ConR1 | | AmpR | Cassette | This work |
| pMM1235 | mScarlet-I | ColE1-CamR-mScarlet-I | | CamR | Part, Type 3 | Harmer et al. 2023 ACS Syn Bio[1] |
| pMM1236 | pGAL1-mScarlet cassette | ConL2-pGAL1-mScarlet-I-tENO1-ConRE | | AmpR | Cassette | Harmer et al. 2023 ACS Syn Bio[1] |
| pMM1245 | eMagB | ColE1-CamR-eMagB | | CamR | Part, Type 3a | Harmer et al. 2023 ACS Syn Bio[1] |
| pMM1248 | pRPL18B-Gal4DBD-eMagA | ConLS-pRPL18B-Gal4DBD-eMagA-tENO1-ConR1 | | AmpR | Cassette | Harmer et al. 2023 ACS Syn Bio[1] |
| pMM1255 | 5xBS CYC180pr-mScarlet-I | ConL2-5xBS CYC180pr-mScarlet-I-tENO1-ConRE | | AmpR | Cassette | This work |
| pMM1256 | pCCW12-NL | 5' Ura3 homology, | URA3 | KanR | Multigene | This work |

| | | | | | | |
|---|---|---|---|---|---|---|
| | S-VP16-EL222-tENO1, 5xBS CYC180pr-mScarlet-I | pCCW12-NLS-VP16-EL222-tENO1, spacer, 5xBS CYC180pr-mScarlet-I-tENO1, Ura3, Ura3 3' homology KanR-ColE1 | | | | |
| pMM1259 | eMagAF | ColE1-CamR-eMagAF | | CamR | Part, Type 3b | Harmer et al. 2023 ACS Syn Bio[1] |
| pMM1263 | pRPL18B-Gal4DBD-eMagAF | ConLS-pRPL18B-Gal4DBD-eMagAF-tENO1-ConR1 | | AmpR | Cassette | Harmer et al. 2023 ACS Syn Bio[1] |
| pMM1264 | pRPL18B-eMagBF-Gal4AD | ConL1-pRPL18B-eMagBF-Gal4AD-tENO1-ConR2 | | AmpR | Cassette | Harmer et al. 2023 ACS Syn Bio[1] |
| pMM1265 | pRPL18B-eMagBM-Gal4AD | ConL1-pRPL18B-eMagBM-Gal4AD-tENO1-ConR2 | | AmpR | Cassette | Harmer et al. 2023 ACS Syn Bio[1] |
| pMM1269 | pRPL18B-Gal4DBD-eMagAF-tENO1, pRPL18B-eMagBF-Gal4AD-tENO1, pGAL1-mScarlet | 5' Ura3 homology, pRPL18B-Gal4DBD-eMagAF-tENO1, pRPL18B-eMagBF-Gal4AD-tENO1, pGAL1-mScarlet-I-tENO1, Ura3, Ura 3' homology KanR-ColE1 | URA3 | KanR | Multigene | Harmer et al. 2023 ACS Syn Bio[1] |
| pMM1270 | pRPL18B-Gal4DBD-eMagA-tENO1, pRPL18B-eMagBM-Gal4AD-tENO1, pGAL1-mScarlet-I | 5' Ura3 homology, pRPL18B-Gal4DBD-eMagA-tENO1, pRPL18B-eMagBM-Gal4AD-tENO1, pGAL1-mScarlet-I-tENO1, Ura3, Ura 3' homology KanR-ColE1 | URA3 | KanR | Multigene | Harmer et al. 2023 ACS Syn Bio[1] |
| pMM1327 | pRPL18B-Gal4DBD-eMagAF-tENO1, | 5' Ura3 homology, pRPL18B-Gal4DBD-eMagAF-tENO1, | URA3 | KanR | Multigene | This work |

| | | | | | | |
|---|---|---|---|---|---|---|
| | pRPL18B-eMagBM-Gal4AD-tENO1, pGAL1-mScarlet-I | pRPL18B-eMagBM-Gal4AD-tENO1, pGAL1-mScarlet-I-tENO1, Ura3, Ura 3' homology KanR-ColE1 | | | | |
| pMM1328 | pRPL18B-Gal4DBD-eMagAF-tENO1, pRPL18B-eMagB-Gal4AD-tENO1, pGAL1-mScarlet | 5' Ura3 homology, pRPL18B-Gal4DBD-eMagAF-tENO1, pRPL18B-eMagB-Gal4AD-tENO1, pGAL1-mScarlet-I-tENO1, Ura3, Ura 3' homology KanR-ColE1 | URA3 | KanR | Multigene | Harmer et al. 2023 ACS Syn Bio[1] |
| pMM1330 | pRPL18B-Gal4DBD-CRY2PHRW349R-tENO1, pRPL18B-Gal4AD-CIB1-tENO1, pGAL1-mScarlet-I | 5' Ura3 homology, pRPL18B-Gal4DBD-CRY2PHRW349R-tENO1, pRPL18B-Gal4AD-CIB1-tENO1, pGAL1-mScarlet-I-tENO1, Ura3, Ura 3' homology KanR-ColE1 | URA3 | KanR | Multigene | This work |
| pMM1331 | pRPL18B-Gal4DBD-CRY2PHRL348F-tENO1, pRPL18B-Gal4AD-CIB1-tENO1, pGAL1-mScarlet-I | 5' Ura3 homology, pRPL18B-Gal4DBD-CRY2PHRL348F-tENO1, pRPL18B-Gal4AD-CIB1-tENO1, pGAL1-mScarlet-I-tENO1, Ura3, Ura 3' homology KanR-ColE1 | URA3 | KanR | Multigene | This work |
| pMM1419 | pLexA-miRFP680 | ConL2-pLexA-miRFP680-tENO1-ConRE | | AmpR | Cassette | This work |
| pMM1430 | Leu2 homology, GFP dropout | 5' Leu2 homology, GFP dropout, Leu2, Leu2 3' homology KanR-ColE1 | LEU2 | KanR | Multigene | This work |
| pMM1431 | NLS-VP16-EL222(A79Q) | ColE1-CamR-NLS-VP16-EL222(A79Q) | | CamR | Part, Type 3 | This work |
| pMM1432 | NLS-VP16-E | ColE1-CamR-NLS-V | | CamR | Part, | This work |

| | | | | | | |
|---|---|---|---|---|---|---|
| | L222 AQTrip | P16-EL222 AQTrip | | | Type 3 | |
| pMM1435 | p65AD | ColE1-CamR-p65AD | | CamR | Part, Type 3b | This work |
| pMM1446 | Msn2AD | ColE1-CamR-Msn2AD | | CamR | Part, Type 3b | This work |
| pMM1447 | VP16 | ColE1-CamR-VP16 | | CamR | Part, Type 3b | This work |
| pMM1452 | pCCW12-NLS-VP16-EL222(A79Q) | ConLS-pCCW12-NLS-VP16-EL222(A79Q)-tENO1-ConR1 | | AmpR | Cassette | This work |
| pMM1453 | pCCW12-NLS-VP16-EL222AQTrip | ConLS-pCCW12-NLS-VP16-EL222AQTrip-tENO1-ConR1 | | AmpR | Cassette | This work |
| pMM1458 | pRPL18B-eMagB-p65AD | ConL1-pRPL18B-eMagB-p65AD-tENO1-ConR2 | | AmpR | Cassette | This work |
| pMM1459 | pRPL18B-eMagB-Msn2AD | ConL1-pRPL18B-eMagB-Msn2AD-tENO1-ConR2 | | AmpR | Cassette | This work |
| pMM1460 | pRPL18B-eMagB-VP16AD | ConL1-pRPL18B-eMagB-VP16AD-tENO1-ConR2 | | AmpR | Cassette | This work |
| pMM1469 | pRPL18B-LexA-eMagAF | ConLS-pRPL18B-LexA-eMagAF-tENO1-ConR1 | | AmpR | Cassette | This work |
| pMM1473 | pCCW12-NLS-VP16-EL222(A79Q)-tENO1, spacer, 5xBS CYC180pr-mScarlet-I | 5' Ura3 homology, pCCW12-NLS-VP16-EL222(A79Q)-tENO1, spacer, 5xBS CYC180pr-mScarlet-I-tENO1, Ura3, Ura 3' homology KanR-ColE1 | URA3 | KanR | Multigene | This work |
| pMM1474 | pCCW12-NLS-VP16-EL222AQTrip-tENO1, spacer, 5xBS CYC180pr-mScarlet-I | 5' Ura3 homology, pCCW12-NLS-VP16-EL222AQTrip-tENO1, spacer, 5xBS CYC180pr-mScarlet-I-tENO1, Ura3, Ura 3' homology | URA3 | KanR | Multigene | This work |

| | | | | | | |
|---|---|---|---|---|---|---|
| | | KanR-ColE1 | | | | |
| pMM1480 | pRPL18B-Gal4DBD-eMagA-tENO1, pRPL18B-eMagB-p65AD-tENO1, pGAL1-mScarlet-I | 5' Ura3 homology, pRPL18B-Gal4DBD-eMagA-tENO1, pRPL18B-eMagB-p65AD-tENO1, pGAL1-mScarlet-I-tENO1, Ura3, Ura 3' homology<br>KanR-ColE1 | URA3 | KanR | Multigene | This work |
| pMM1481 | pRPL18B-Gal4DBD-eMagA-tENO1, pRPL18B-eMagB-Msn2AD-tENO1, pGAL1-mScarlet-I | 5' Ura3 homology, pRPL18B-Gal4DBD-eMagA-tENO1, pRPL18B-eMagB-Msn2AD-tENO1, pGAL1-mScarlet-I-tENO1, Ura3, Ura 3' homology<br>KanR-ColE1 | URA3 | KanR | Multigene | This work |
| pMM1482 | pRPL18B-Gal4DBD-eMagA-tENO1, pRPL18B-eMagB-VP16AD-tENO1, pGAL1-mScarlet-I | 5' Ura3 homology, pRPL18B-Gal4DBD-eMagA-tENO1, pRPL18B-eMagB-VP16AD-tENO1, pGAL1-mScarlet-I-tENO1, Ura3, Ura 3' homology<br>KanR-ColE1 | URA3 | KanR | Multigene | This work |
| pMM1485 | pRPL18B-LexA-eMagAF-tENO1, pRPL18B-eMagBF-Gal4AD-tENO1, pLexA-miRFP680 | 5' Leu2 homology, pRPL18B-LexA-eMagAF-tENO1, pRPL18B-eMagBF-Gal4AD-tENO1, pLexA-miRFP680-tENO1, Leu2, Leu2 3' homology<br>KanR-ColE1 | LEU2 | KanR | Multigene | This work |

#### Table S4: Gene blocks used in this study

| ID | Alias | Sequence | Purpose |
|---|---|---|---|
| gMM041 | miRFP680 | GCATCGTCTCATCGGTCTCATATGGCGGAAGGCT | miRFP680 |

| | | | |
|---|---|---|---|
| | entry part 3 | CCGTCGCCAGGCAGCCTGACCTCTTGACCTGCGACGATGAGCCGATCCATATCCCCGGTGCCATCCAACCGCATGGACTtCTtCTCGCCCTCGCCGCCGACATGACGATCGTTGCCGGCAGCGACAACCTTCCCGAACTCACCGGACTtGCGATCGGCGCCCTtATCGGCCGCTCTGCGGCCGATGTCTTCGACTCGGAAACGCACAACCGTCTtACGATCGCCTTGGCCGAGCCCGGGGCGGCCGTCGGAGCACCGATCACTGTCGGCTTCACGATGCGAAAGGACGCAGGCTTCATCGGCTCCTGGCATCGCCATGATCAGCTCATCTTCCTCGAGCTCGAGCCTCCCCAGCGGGACGTCGCCGAGCCGCAGGCGTTCTTCCGCCGCACCAACAGCGCCATCCGCCGCCTtCAGGCCGCCGAAACCTTGGAAAGCGCCTGCGCCGCCGCGGCGCAAGAGGTGCGGAAGATTACCGGCTTCGATCGGGTGATGATCTATCGCTTCGCCTCCGACTTCAGCGGCGAAGTGATCGCAGAGGATCGGTGCGCCGAGGTCGAGTCAAAACTAGGCCTtCACTATCCTGCCTCAACCGTGCCGGCGCAGGCCCGTCGGCTCTATACCATCAACCCGGTACGGATCATTCCCGATATCAATTATCGGCCGGTGCCGGTCACCCCAGACCTCAATCCGGTCACCGGGCGGCCGATTGATCTTAGCTTCGCCATCCTtCGCAGCGTGTCGCCCTGCCATCTtGAGTTCATGCGCAACATAGGCATGCACGGCACGATGTCGATCTCGATTTTGCGCGGCGAGCGACTtTGGGGATTGATCGTTTGCCATCACCGAACGCCGTACTACGTCGATCTCGATGGCCGCCAAGCCTGCAAGAGGGTCGCCGAGAGGCTtGCCACTCAGATCGGCGTGATGGAAGAGccATCCTGAGACCTGAGACGGCAT | part 3 plasmid for yOTK |
| gMM046 | 5xBS-CYC 180pr fragment | gcatCGTCTCaATAGGTAGCCTTTAGTCCATGAAGCTTAGACACTAGAGGGACTAGAGTGCTGACACTACAGGCATATATATATGTGTGCGACGACACATGATCATATGGCATGCATGTGCTCTGTATGTATATAAAACTCTTGTTTTCTTCTTTTCTCTAAATATTCTTTCCTTATACATTAGGACCTTTGCAGCATAAATTACTATACTTCTATAGACACACAAACACAAATACagatcTATGTGAGACCTGAGACGGCATGC | Makes 5xBS-CYC180pr type 2 part (BsmBI gga) used for pMM1098 |
| gMM054 | NLS-VP16 AD-EL222 | gcatCGTCTCaTCGGTCTCaTATGGGCCCTAAAAAGAAGCGTAAAGTCGCCCCCCCGACCGATGTCAGCCTGGGGGACGAGCTCCACTTAGACGGCGAGGACGTGGCGATGGCGCATGCCGACGCGCTAGACGATTTCGATCTGGACATGTTGGGGGACGGGGATTCCCCGGGTCCGGGATTTACCCCCCACGACTCCGCCCCCTACGGCGCTCTGGATATGGCCGACTTCGAGTTTGAGCAGATGTTTACCGATGCCCTTGGAATTGACGAGTACGGTGGGGAATTCGGGGCAGACGACACACGCGTTGAGGTGCAACCGCCGGCGCAGTGGGTCCTCGACCTGATCGAGGCCAGCCCGATCGCATCG | Makes NLS-VP16AD-EL222 type 3 part |

| | | | |
|---|---|---|---|
| | | GTCGTGTCCGATCCGCGACTCGCCGACAATCCG CTGATCGCCATCAACCAGGCCTTCACCGACCTGA CCGGCTATTCCGAAGAAGAATGCGTCGGCCGCAA TTGCCGATTCCTGGCAGGTTCCGGCACCGAGCC GTGGCTGACCGACAAGATCCGCCAAGGCGTGCG CGAGCACAAGCCGGTGCTGGTCGAGATCCTGAA CTACAAGAAGGACGGCACGCCGTTCCGCAATGC CGTGCTCGTTGCACCGATCTACGATGACGACGAC GAGCTTCTCTATTTCCTCGGCAGCCAGGTCGAAG TCGACGACGACCAGCCCAACATGGGCATGGCGC GCCGCGAACGCGCCGCGGAAATGCTCAAGACGC TGTCGCCGCGCCAGCTCGAGGTTACGACGCTGG TGGCATCGGGCTTGCGCAACAAGGAAGTGGCGG CCCGGCTCGGCCTGTCGGAGAAAACCGTCAAGA TGCACCGCGGGCTGGTGATGGAAAAGCTCAACC TGAAGACCAGTGCCGATCTGGTGCGCATTGCCGT CGAAGCCGGAATCTAAATCCtGAGACCtGAGACGg cat | |
| gMM068 | NLS-VP16AD-EL222(AQTrip) | gcatCGTCTCaTCGGTCTCaTATGGGCCCTAAAAAG AAGCGTAAAGTCGCCCCCCCGACCGATGTCAGC CTGGGGGACGAGCTCCACTTAGACGGCGAGGAC GTGGCGATGGCGCATGCCGACGCGCTAGACGAT TTCGATCTGGACATGTTGGGGGACGGGGATTCCC CGGGTCCGGGATTTACCCCCCACGACTCCGCCC CCTACGGCGCTCTGGATATGGCCGACTTCGAGTT TGAGCAGATGTTTACCGATGCCCTTGGAATTGAC GAGTACGGTGGGGAATTCGGGGCAGACGACACA CGCGTTGAGGTGCAACCGCCGGCGCAGTGGGTC CTCGACCTGATCGAGGCCAGCCCGATCGCATCGA TTGTGTCCGATCCGCGACTCGCCGACAATCCGAT TATCGCCATCAACCAGGCCTTCACCGACCTGACC GGCTATTCCGAAGAAGAATGCGTCGGCCGCAATT GCCGATTCCTGCAAGGTTCCGGCACCGAGCCGT GGCTGACCGACAAGATCCGCCAAGGCGTGCGCG AGCACAAGCCGGTGCTGGTCGAGATCCTGAACTA CAAGAAGGACGGCACGCCGTTCCGCAATGCCGT GCTCATTGCACCGATCTACGATGACGACGACGAG CTTCTCTATTTCCTCGGCAGCCAGGTCGAAGTCG ACGACGACCAGCCCAACATGGGCATGGCGCGCC GCGAACGCGCCGCGGAAATGCTCAAGACGCTGT CGCCGCGCCAGCTCGAGGTTACGACGCTGGTGG CATCGGGCTTGCGCAACAAGGAAGTGGCGGCCC GGCTCGGCCTGTCGGAGAAAACCGTCAAGATGC ACCGCGGGCTGGTGATGGAAAAGCTCAACCTGA AGACCAGTGCCGATCTGGTGCGCATTGCCGTCGA AGCCGGAATCTAAATCCtGAGACCtGAGACGgcat | Makes NLS-VP16AD-EL222(AQTrip) type 3 part |
| gMM071 | p65AD entry part | gcatCGTCTCaTCGGTCTCAttctgaattccaataccttccaga cacggatgatcgccatcgaatcgaagagaagagaaaacgcacctac | Makes p65AD type |

| | | | |
|---|---|---|---|
| | 3b | gagacgttcaaatctattatgaagaaatctcccttcagtgggccgacggacccaaggccgccaccgcgaaggatagccgttccatcaagaagctcagcttctgtacccaaacccgccccacagccttacccctttacttcctccctctccactatcaactacgatgagttccccacgatggtttttccttcaggacagatatcccaagcgagcgccctcgcaccagccccaccacaagtgcttcctcaggcccctgcgcctgctccggcaccggcgatggtgagcgctctggctcaagctcccgcgccagtccctgttttggcaccagggccacctcaggcagtggctccgcccgctccaaaacctactcaagcgggcgaaggaactctgagcgaggcgctcctgcagcttcaatttgatgacgaagatctcggcgcactcctcggtaattcaacggaccccgctgttttcactgacctggcaagcgtggataactctgaattccaacagctccttaaccagggcataccggtcgcgcctcatacaactgaaccaatgctgatggaatatccggaggcaataaccagacttgtgacgggggcgcagcgaccgcctgatccagcacccgcaccgcttggggcgcctggcttgcccaatggactcctttctggcgacgaggacttttccagcatcgcagacatggacttttctgcactcctttctcagatttcctcaatcctGAGACCtGAGACGgcat | 3b part |

### Table S5: Sequences used in this study

| Construct | Sequence |
|---|---|
| Gal4DBD-CRY2 (535) | ATGaagctactgtcttctatcgaacaagcatgcgatatttgccgacttaaaaagctcaagtgctccaaagaaaaaccgaagtgcgccaagtgtctgaagaacaactgggagtgtcgctactctcccaaaaccaaaaggtcaccgctgactagggcacatctgacagaagtggaatcaaggctagaaagactggaacagctatttctactgatttttcctcgagaagaccttgacatgattttgaaaatggattctttacaggatataaaagcattgttaacaggattatttgtacaagataatgtgaataaagatgccgtcacagatagattggcttcagtggagactgatatgcctctaacattgagacagcatagaataagtgcgacatcatcatcggaagagagtagtaacaaaggtcaaagacagttgactgtatcgggTTCTacaggtgctagcttcatgaagatggacaaaaagactatagtttggtttagaagagatctaaggattgaggataatcctgcattagcagcagctgctcacgaaggatctgtttttcctgtcttcatttggtgtcctgaagaagaaggacagttttatcctggaagagcttcaagatggtggatgaaacaatcacttgctcacttatctcaatccttgaaggctcttggatctgacctcactttaatcaaaacccacaacacgatttcagcgatcttggattgtatccgcgttaccggtgctacaaaagtcgtctttaaccacctctatgatcctgtttcgttagttcgggaccataccgtaaaggagaagctggtggaacgtgggatctctgtgcaaagctacaatggagatctattgtatgaaccgtgggagatatactgcgaaaagggcaaacctttacgagtttcaattcttactggaagaaatgcttagatatgtcgattgaatccgttatgcttcctccttggcggttgatgccaataactgcagcggctgaagcgatttgggcgtgttcgattgaagaactagggctggagaatgaggccgagaaaccgagcaatgcgttgttaactagagcttggagtccaggatggagcaatgctgataagttactaaatgagttcatcgagaagcagttgatagattatgcaaagaacagcaagaaagttgttgggaattctacttcactactttctccgtatctccatttcggggaaataagcgtcagacacgttttccagtgtgcccggatgaaacaaattatatgggcaagagataagaacagtgaaggagaagaaagtgcagatcttttttcttaggggaatcggtttaagagagtattctcggtatatatgtttcaacttcccgtttactcacgagcaatcgttgttgagtcatcttcggtttttcccttgggatgctgatgttgataagttcaaggcctggagacaaggcaggaccggttatccgttggtggatgccggaatgagagagctttgggctaccggatggatgcataacagaataagagtgattgtttcaagctttgctgtgaagtttcttctccttccatggaaatggggaatgaagtatttctgggatacactttggatgctgatttggaatgtgacatccttggctggcagtatatctctgggagtatccccgatggccacgagcttgatcgcttggacaatcccgcgttacaaggcgccaaatatgacccagaaggtgagtac |

| | |
|---|---|
| | ataaggcaatggcttcccgagcttgcgagattgccaactgaatggatccatcatccatgggacgctcctt<br>taaccgtactcaaagcttctggtgtggaactcggaacaaactatgcgaaacccattgtagacatcgac<br>acagctcgtgagctactagctaaagctatttcaagaacccgtgaagcacagatcatgatcggagcag<br>cacctgatgagattgtagcagatagcttcgaggccttaggggctaataccattaaagaacctggtctttg<br>cccatctgtgtcttctaatgaccaacaagtaccttcggctgttGGatcctaa |
| Gal4DBD-CRY2 PHR | ATGaagctactgtcttctatcgaacaagcatgcgatatttgccgacttaaaaagctcaagtgctccaaa<br>gaaaaaccgaagtgcgccaagtgtctgaagaacaactgggagtgtcgctactctcccaaaaccaaa<br>aggtcaccgctgactagggcacatctgacagaagtggaatcaaggctagaaagactggaacagcta<br>tttctactgatttttcctcgagaagaccttgacatgattttgaaaatggattctttacaggatataaaagcatt<br>gttaacaggattatttgtacaagataatgtgaataaagatgccgtcacagatagattggcttcagtggag<br>actgatatgcctctaacattgagacagcatagaataagtgcgacatcatcatcggaagagagtagtaa<br>caaaggtcaaagacagttgactgtatcgggTTCTacaggtgctagcttcatgaagatggacaaaaa<br>gactatagtttggtttagaagagatctaaggattgaggataatcctgcattagcagcagctgctcacgaa<br>ggatctgtttttcctgtcttcatttggtgtcctgaagaagaaggacagttttatcctggaagagcttcaagat<br>ggtggatgaaacaatcacttgctcacttatctcaatccttgaaggctcttggatctgacctcactttaatca<br>aaacccacaacacgatttcagcgatcttggattgtatccgcgttaccggtgctacaaaagtcgtctttaac<br>cacctctatgatcctgtttcgttagttcgggaccataccgtaaaggagaagctggtggaacgtgggatct<br>ctgtgcaaagctacaatggagatctattgtatgaaccgtgggagatatactgcgaaaagggcaaacct<br>tttacgagtttcaattcttactggaagaaatgcttagatatgtcgattgaatccgttatgcttcctcctccttgg<br>cggttgatgccaataactgcagcggctgaagcgatttgggcgtgttcgattgaagaactagggctgga<br>gaatgaggccgagaaaccgagcaatgcgttgttaactagagcttggagtccaggatggagcaatgct<br>gataagttactaaatgagttcatcgagaagcagttgatagattatgcaaagaacagcaagaaagttgtt<br>gggaattctacttcactactttctccgtatctccatttcggggaaataagcgtcagacacgttttccagtgtg<br>cccggatgaaacaaattatatgggcaagagataagaacagtgaaggagaagaaagtgcagatcttt<br>ttcttaggggaatcggtttaagagagtattctcggtatatatgtttcaacttcccgtttactcacgagcaatcg<br>ttgttgagtcatcttcggtttttcccttgggatgctgatgttgataagttcaaggcctggagacaaggcagg<br>accggttatccgttggtggatgccggaatgagagagctttgggctaccggatggatgcataacagaat<br>aagagtgattgtttcaagctttgctgtgaagtttcttctccttccatggaaatggggaatgaagtatttctggg<br>atacactttggatgctgatttggaatgtgacatccttggctggcagtatatctctgggagtatccccgatgg<br>ccacgagcttgatcgcttggacaatcccgcgttacaaggcgccaaatatgacccagaaggtgagtac<br>ataaggcaatggcttcccgagcttgcgagattgccaactgaatggatccatcatccatgggacgctcctt<br>taaccgtactcaaagcttctggtgtggaactcggaacaaactatgcgaaacccattgtagacatcgac<br>acagctcgtgagctactagctaaagctatttcaagaacccgtgaagcacagatcatgatcggagcag<br>caGCCCGGGGatcctaa |
| Gal4DBD-CRY2 PHR(W349R) | ATGaagctactgtcttctatcgaacaagcatgcgatatttgccgacttaaaaagctcaagtgctccaaa<br>gaaaaaccgaagtgcgccaagtgtctgaagaacaactgggagtgtcgctactctcccaaaaccaaa<br>aggtcaccgctgactagggcacatctgacagaagtggaatcaaggctagaaagactggaacagcta<br>tttctactgatttttcctcgagaagaccttgacatgattttgaaaatggattctttacaggatataaaagcatt<br>gttaacaggattatttgtacaagataatgtgaataaagatgccgtcacagatagattggcttcagtggag<br>actgatatgcctctaacattgagacagcatagaataagtgcgacatcatcatcggaagagagtagtaa<br>caaaggtcaaagacagttgactgtatcgggTTCTacaggtgctagcttcatgaagatggacaaaaa<br>gactatagtttggtttagaagagatctaaggattgaggataatcctgcattagcagcagctgctcacgaa<br>ggatctgtttttcctgtcttcatttggtgtcctgaagaagaaggacagttttatcctggaagagcttcaagat<br>ggtggatgaaacaatcacttgctcacttatctcaatccttgaaggctcttggatctgacctcactttaatca<br>aaacccacaacacgatttcagcgatcttggattgtatccgcgttaccggtgctacaaaagtcgtctttaac<br>cacctctatgatcctgtttcgttagttcgggaccataccgtaaaggagaagctggtggaacgtgggatct<br>ctgtgcaaagctacaatggagatctattgtatgaaccgtgggagatatactgcgaaaagggcaaacct<br>tttacgagtttcaattcttactggaagaaatgcttagatatgtcgattgaatccgttatgcttcctcctccttgg |

| | |
|---|---|
| | cggttgatgccaataactgcagcggctgaagcgatttgggcgtgttcgattgaagaactagggctgga<br>gaatgaggccgagaaaccgagcaatgcgttgttaactagagcttggagtccaggatggagcaatgct<br>gataagttactaaatgagttcatcgagaagcagttgatagattatgcaaagaacagcaagaaagttgtt<br>gggaattctacttcactactttctccgtatctccatttcggggaaataagcgtcagacacgttttccagtgtg<br>cccggatgaaacaaattatatgggcaagagataagaacagtgaaggagaagaaagtgcagatcttt<br>ttcttaggggaatcggtttaagagagtattctcggtatatatgtttcaacttcccgtttactcacgagcaatcg<br>ttgttgagtcatcttcggtttttcccttgggatgctgatgttgataagttcaaggcctggagacaaggcagg<br>accggttatccgttggtggatgccggaatgagagagcttagggctaccggatggatgcataacagaat<br>aagagtgattgtttcaagctttgctgtgaagtttcttctccttccatggaaatggggaatgaagtatttctggg<br>atacacttttggatgctgatttggaatgtgacatccttggctggcagtatatctctgggagtatccccgatgg<br>ccacgagcttgatcgcttggacaatcccgcgttacaaggcgccaaatatgacccagaaggtgagtac<br>ataaggcaatggcttcccgagcttgcgagattgccaactgaatggatccatcatccatgggacgctcctt<br>taaccgtactcaaagcttctggtgtggaactcggaacaaactatgcgaaacccattgtagacatcgac<br>acagctcgtgagctactagctaaagctatttcaagaacccgtgaagcacagatcatgatcggagcag<br>caGCCCGGGGatcctaa |
| Gal4DBD-CRY2 PHR(L348F) | ATGaagctactgtcttctatcgaacaagcatgcgatatttgccgacttaaaaagctcaagtgctccaaa<br>gaaaaaccgaagtgcgccaagtgtctgaagaacaactgggagtgtcgctactctcccaaaaccaaa<br>aggtcaccgctgactagggcacatctgacagaagtggaatcaaggctagaaagactggaacagcta<br>tttctactgatttttcctcgagaagaccttgacatgattttgaaaatggattctttacaggatataaaagcatt<br>gttaacaggattatttgtacaagataatgtgaataaagatgccgtcacagatagattggcttcagtggag<br>actgatatgcctctaacattgagacagcatagaataagtgcgacatcatcatcggaagagagtagtaa<br>caaaggtcaaagacagttgactgtatcgggTTCTacaggtgctagcttcatgaagatggacaaaaa<br>gactatagtttggtttagaagagatctaaggattgaggataatcctgcattagcagcagctgctcacgaa<br>ggatctgtttttcctgtcttcatttggtgtcctgaagaagaaggacagttttatcctggaagagcttcaagat<br>ggtggatgaaacaatcacttgctcacttatctcaatccttgaaggctcttggatctgacctcactttaatca<br>aaacccacaacacgatttcagcgatcttggattgtatccgcgttaccggtgctacaaaagtcgtctttaac<br>cacctctatgatcctgtttcgttagttcgggaccataccgtaaaggagaagctggtggaacgtgggatct<br>ctgtgcaaagctacaatggagatctattgtatgaaccgtgggagatatactgcgaaaagggcaaacct<br>tttacgagtttcaattcttactggaagaaatgcttagatatgtcgattgaatccgttatgcttcctcctccttgg<br>cggttgatgccaataactgcagcggctgaagcgatttgggcgtgttcgattgaagaactagggctgga<br>gaatgaggccgagaaaccgagcaatgcgttgttaactagagcttggagtccaggatggagcaatgct<br>gataagttactaaatgagttcatcgagaagcagttgatagattatgcaaagaacagcaagaaagttgtt<br>gggaattctacttcactactttctccgtatctccatttcggggaaataagcgtcagacacgttttccagtgtg<br>cccggatgaaacaaattatatgggcaagagataagaacagtgaaggagaagaaagtgcagatcttt<br>ttcttaggggaatcggtttaagagagtattctcggtatatatgtttcaacttcccgtttactcacgagcaatcg<br>ttgttgagtcatcttcggtttttcccttgggatgctgatgttgataagttcaaggcctggagacaaggcagg<br>accggttatccgttggtggatgccggaatgagagagttttgggctaccggatggatgcataacagaata<br>agagtgattgtttcaagctttgctgtgaagtttcttctccttccatggaaatggggaatgaagtatttctggga<br>tacacttttggatgctgatttggaatgtgacatccttggctggcagtatatctctgggagtatccccgatggc<br>cacgagcttgatcgcttggacaatcccgcgttacaaggcgccaaatatgacccagaaggtgagtaca<br>taaggcaatggcttcccgagcttgcgagattgccaactgaatggatccatcatccatgggacgctccttt<br>aaccgtactcaaagcttctggtgtggaactcggaacaaactatgcgaaacccattgtagacatcgaca<br>cagctcgtgagctactagctaaagctatttcaagaacccgtgaagcacagatcatgatcggagcagc<br>aGCCCGGGGatcctaa |
| Gal4DBD-CRY2 FL | ATGaagctactgtcttctatcgaacaagcatgcgatatttgccgacttaaaaagctcaagtgctccaaa<br>gaaaaaccgaagtgcgccaagtgtctgaagaacaactgggagtgtcgctactctcccaaaaccaaa<br>aggtcaccgctgactagggcacatctgacagaagtggaatcaaggctagaaagactggaacagcta<br>tttctactgatttttcctcgagaagaccttgacatgattttgaaaatggattctttacaggatataaaagcatt |

| | |
|---|---|
| | gttaacaggattatttgtacaagataatgtgaataaagatgccgtcacagatagattggcttcagtggag<br>actgatatgcctctaacattgagacagcatagaataagtgcgacatcatcatcggaagagagtagtaa<br>caaaggtcaaagacagttgactgtatcgggTTCTacaggtgctagcttcatgaagatggacaaaaa<br>gactatagtttggtttagaagagatctaaggattgaggataatcctgcattagcagcagctgctcacgaa<br>ggatctgtttttcctgtcttcatttggtgtcctgaagaagaaggacagttttatcctggaagagcttcaagat<br>ggtggatgaaacaatcacttgctcacttatctcaatccttgaaggctcttggatctgacctcactttaatca<br>aaacccacaacacgatttcagcgatcttggattgtatccgcgttaccggtgctacaaaagtcgtctttaac<br>cacctctatgatcctgtttcgttagttcgggaccataccgtaaaggagaagctggtggaacgtgggatct<br>ctgtgcaaagctacaatggagatctattgtatgaaccgtgggagatatactgcgaaaagggcaaacct<br>tttacgagtttcaattcttactggaagaaatgcttagatatgtcgattgaatccgttatgcttcctcctccttgg<br>cggttgatgccaataactgcagcggctgaagcgatttgggcgtgttcgattgaagaactagggctgga<br>gaatgaggccgagaaaccgagcaatgcgttgttaactagagcttggagtccaggatggagcaatgct<br>gataagttactaaatgagttcatcgagaagcagttgatagattatgcaaagaacagcaagaaagttgtt<br>gggaattctacttcactactttctccgtatctccatttcggggaaataagcgtcagacacgttttccagtgtg<br>cccggatgaaacaaattatatgggcaagagataagaacagtgaaggagaagaaagtgcagatcttt<br>ttcttaggggaatcggtttaagagagtattctcggtatatatgtttcaacttcccgtttactcacgagcaatcg<br>ttgttgagtcatcttcggtttttcccttgggatgctgatgttgataagttcaaggcctggagacaaggcagg<br>accggttatccgttggtggatgccggaatgagagagctttgggctaccggatggatgcataacagaat<br>aagagtgattgtttcaagctttgctgtgaagtttcttctccttccatggaaatggggaatgaagtatttctggg<br>atacacttttggatgctgatttggaatgtgacatccttggctggcagtatatctctgggagtatccccgatgg<br>ccacgagcttgatcgcttggacaatcccgcgttacaaggcgccaaatatgacccagaaggtgagtac<br>ataaggcaatggcttcccgagcttgcgagattgccaactgaatggatccatcatccatgggacgctcctt<br>taaccgtactcaaagcttctggtgtggaactcggaacaaactatgcgaaacccattgtagacatcgac<br>acagctcgtgagctactagctaaagctatttcaagaacccgtgaagcacagatcatgatcggagcag<br>cacctgatgagattgtagcagatagcttcgaggccttaggggctaataccattaaagaacctggtctttg<br>cccatctgtgtcttctaatgaccaacaagtaccttcggctgttcgttacaacgggtcaaagagagtgaaa<br>cctgaggaagaagaagagagagacatgaagaaatctaggggattcgatgaaagggagttgttttcg<br>actgctgaatcttcttcttcttcgagtgtgtttttcgtttcgcagtcttgctcgttggcatcagaagggaagaat<br>ctggaaggtattcaagattcatctgatcagattactacaagtttgggaaaaaatggttgcaaaGGatcc<br>taa |
| Gal4AD-CIB1 | atggataaagcggaattaattcccgagcctccaaaaaagaagagaaaggtcgaattgggtaccgcc<br>gccaattttaatcaaagtgggaatattgctgatagctcattgtccttcactttcactaacagtagcaacggt<br>ccgaacctcataacaactcaaacaaattctcaagcgctttcacaaccaattgcctcctctaacgttcatg<br>ataacttcatgaataatgaaatcacggctagtaaaattgatgatggtaataattcaaaaccactgtcacc<br>tggttggacggaccaaactgcgtataacgcgtttggaatcactacagggatgtttaataccactacaatg<br>gatgatgtatataactatctattcgatgatgaagataccccaccaaacccaaaaaaagagatctttaat<br>acgactcactatagggcgagcgccgaagctagcgccaccatgaatggagctataggaggtgaccttt<br>tgctcaattttcctgacatgtcggtcctagagcgccaaagggctcacctcaagtacctcaatcccaccttt<br>gattctcctctcgccggcttctttgccgattcttcaatgattaccggcggcgagatggacagctatctttcga<br>ctgccggtttgaatcttccgatgatgtacggtgagacaacggtggaaggtgattcaagactctcaatttcg<br>ccggaaacgacgcttgggactggaaatttcaagaaacggaagtttgatacagagactaaggattgta<br>atgagaagaagaagaagatgacgatgaacagagatgacctagtagaagaaggagaagaagaga<br>agtcgaaaataacagagcaaaacaatgggagcacaaaaagcatcaagaagatgaaacacaaag<br>ccaagaaagaagagaacaatttctctaatgattcatctaaagtgacgaaggaattggagaaaacgga<br>ttatattcatgttcgtgcacgacgaggccaagccactgatagtcacagcatagcagaacgagttagaa<br>gagaaaagatcagtgagagaatgaagtttctacaagatttggttcctggatgcgacaagatcacaggc<br>aaagcagggatgcttgatgaaatcattaactatgttcagtctcttcagagacaaatcgagttcttatcgat<br>gaaactagcaattgtgaatccaaggccggattttgatatggatgacatttttgccaaagaggttgcctca<br>actccaatgactgtggtgccatctcctgaaatggttctttccggttattctcatgagatggttcactctggttat |

| | |
|---|---|
| | tctagtgagatggttaactccggttaccttcatgtcaatccaatgcagcaagtgaataccagttctgatcc<br>attgtcatgcttcaacaatggcgaagctccttcgatgtgggactctcatgtgcagaatctctatggcaattt<br>aggagtaccggtcatcgagctcgagctgcagatgaatcgtagatacggatcctaa |
| Gal4DBD-eMag A | ATGaagctactgtcttctatcgaacaagcatgcgatatttgccgacttaaaaagctcaagtgctccaaa<br>gaaaaaccgaagtgcgccaagtgtctgaagaacaactgggagtgtcgctactctcccaaaaccaaa<br>aggtcaccgctgactagggcacatctgacagaagtggaatcaaggctagaaagactggaacagcta<br>tttctactgatttttcctcgagaagaccttgacatgattttgaaaatggattctttacaggatataaaagcatt<br>gttaacaggattatttgtacaagataatgtgaataaagatgccgtcacagatagattggcttcagtggag<br>actgatatgcctctaacattgagacagcatagaataagtgcgacatcatcatcggaagagagtagtaa<br>caaaggtcaaagacagttgactgtatcgggttctggaggcggaggctccggtggtggtggaagtggc<br>ggtggcggatccatgggacacactctttacgcccctggaggatacgacattatgggatatttggatcag<br>attgcgaaccgcccaaaccctcaggtcgaactggggcctgtggacctgtcatgtgccctgatcctgtgc<br>gatctgaagcaaaaggacactccgatcgtctacgcctcggaagccttcttggagatgaccggataca<br>acagacatgaggtgctcggcaggaactgcagattcctgcagtcccccgacgggatggtgaaaccaa<br>agtcgactcgcaaatatgtggactcgaacacgatctacaccatcaagaaggccatcgaccggaacg<br>ccgaggtccaggtggaggtggtcaactttaagaagaacggccagcggttcgtgaactttctgaccatc<br>attccggtccgggatgaaaccggagagtacagatactccatcggattccagtgcgaaaccgaaTAA |
| eMagB-Gal4AD | atgggacataccctctacgcgccgggggttatgacatcatgggttacctcagacagatcagaaacc<br>ggccgaacccacaagtggagctgggacccgtcgacctctcctgcgccctcgtgctgtgtgaccttaag<br>cagaaggacacccctgtggtgtacgcctccgaagcattcctggagatgaccgggtacaacagacac<br>gaagtgctgggacggaactgccgcttcctgcaatccccggatggaatggtgaagcctaagtcaaccc<br>gcaaatacgtggactccaacactatctacaccatgaagaaggccattgaccgcaatgctgaggtgca<br>agtggaagtggtgaacttcaagaagaacggacagcgcttcgtcaacttcctgactatgattcccgtgcg<br>ggacgaaaccggcgaataccggtacagcatcgggtttcagtgcgagactgagGGCGGTGGC<br>GGCAGTGGTGGAATGAAGCAACTCGAGGACAAGGTTGAGGAACTGC<br>TGAGTAAGAATTACCACCTCGAAAACGAGGTCGCACGATTGAAAAAGT<br>TGGTGGGTGAGggaggtggtggatcgggtggaggTTCTGATAAAGCGGAATTAA<br>TTCCCGAGCCTCCAAAAAAGAAGAGAAAGGTCGAATTGGGTACCGCC<br>GCCAATTTTAATCAAAGTGGGAATATTGCTGATAGCTCATTGTCCTTCA<br>CTTTCACTAACAGTAGCAACGGTCCGAACCTCATAACAACTCAAACAA<br>ATTCTCAAGCGCTTTCACAACCAATTGCCTCCTCTAACGTTCATGATAA<br>CTTCATGAATAATGAAATCACGGCTAGTAAAATTGATGATGGTAATAATT<br>CAAAACCACTGTCACCTGGTTGGACGGACCAAACTGCGTATAACGCG<br>TTTGGAATCACTACAGGGATGTTTAATACCACTACAATGGATGATGTATA<br>TAACTATCTATTCGATGATGAAGATACCCCACCAAACCCAAAAAAAGAGt<br>aa |
| Gal4DBD-eMag AF | ATGaagctactgtcttctatcgaacaagcatgcgatatttgccgacttaaaaagctcaagtgctccaaa<br>gaaaaaccgaagtgcgccaagtgtctgaagaacaactgggagtgtcgctactctcccaaaaccaaa<br>aggtcaccgctgactagggcacatctgacagaagtggaatcaaggctagaaagactggaacagcta<br>tttctactgatttttcctcgagaagaccttgacatgattttgaaaatggattctttacaggatataaaagcatt<br>gttaacaggattatttgtacaagataatgtgaataaagatgccgtcacagatagattggcttcagtggag<br>actgatatgcctctaacattgagacagcatagaataagtgcgacatcatcatcggaagagagtagtaa<br>caaaggtcaaagacagttgactgtatcgggttctggaggcggaggctccggtggtggtggaagtggc<br>ggtggcggatccatgggacacactctttacgcccctggaggatacgacattatgggatatttggatcag<br>attgcgaaccgcccaaaccctcaggtcgaactggggcctgtggacctgtcatgtgccctgatcctgtgc<br>gatctgaagcaaaaggacactccgatcgtctacgcctcggaagccttcttggagatgaccggataca<br>acagacatgaggtgctcggcaggaactgcagattcctgcagtcccccgacgggatggtgaaaccaa |

| | |
|---|---|
| | agtcgactcgcaaatatgtggactcgaacacgatcttcaccatcaagaaggccatcgaccggaacgccgaggtccaggtggaggtggtcaactttaagaagaacggccagcggttcgtgaactttctgaccatcattccggtccgggatgaaaccggagagtacagatactccatcggattccagtgcgaaaccgaaTAA |
| eMagBF-Gal4AD | atgggacataccctctacgcgccggggggttatgacatcatgggttacctcagacagatcagaaaccggccgaacccacaagtggagctgggacccgtcgacctctcctgcgccctcgtgctgtgtgaccttaagcagaaggacacccctgtggtgtacgcctccgaagcattcctggagatgaccgggtacaacagacacgaagtgctgggacggaactgccgcttcctgcaatccccggatggaatggtgaagcctaagtcaacccgcaaatacgtggactccaacactatcttcaccatgaagaaggccattgaccgcaatgctgaggtgcaagtggaagtggtgaacttcaagaagaacggacagcgcttcgtcaacttcctgactatgattcccgtgcgggacgaaaccggcgaataccggtacagcatcgggtttcagtgcgagactgagGGCGGTGGCGGCAGTGGTGGAATGAAGCAACTCGAGGACAAGGTTGAGGAACTGCTGAGTAAGAATTACCACCTCGAAAACGAGGTCGCACGATTGAAAAAGTTGGTGGGTGAGggaggtggtggatcgggtggaggTTCTGATAAAGCGGAATTAATTCCCGAGCCTCCAAAAAAGAAGAGAAAGGTCGAATTGGGTACCGCCGCCAATTTTAATCAAAGTGGGAATATTGCTGATAGCTCATTGTCCTTCACTTTCACTAACAGTAGCAACGGTCCGAACCTCATAACAACTCAAACAAATTCTCAAGCGCTTTCACAACCAATTGCCTCCTCTAACGTTCATGATAACTTCATGAATAATGAAATCACGGCTAGTAAAATTGATGATGGTAATAATTCAAAACCACTGTCACCTGGTTGGACGGACCAAACTGCGTATAACGCGTTTGGAATCACTACAGGGATGTTTAATACCACTACAATGGATGATGTATATAACTATCTATTCGATGATGAAGATACCCCACCAAACCCAAAAAAAGAGtaa |
| eMagBM-Gal4AD | atgggacataccctctacgcgccggggggttatgacatcatgggttacctcagacagatcagaaaccggccgaacccacaagtggagctgggacccgtcgacctctcctgcgccctcatcctgtgtgaccttaagcagaaggacacccctgtggtgtacgcctccgaagcattcctggagatgaccgggtacaacagacacgaagtgctgggacggaactgccgcttcctgcaatccccggatggaatggtgaagcctaagtcaacccgcaaatacgtggactccaacactatctacaccatgaagaaggccattgaccgcaatgctgaggtgcaagtggaagtggtgaacttcaagaagaacggacagcgcttcgtcaacttcctgactatgattcccgtgcgggacgaaaccggcgaataccggtacagcatcgggtttcagtgcgagactgagGGCGGTGGCGGCAGTGGTGGAATGAAGCAACTCGAGGACAAGGTTGAGGAACTGCTGAGTAAGAATTACCACCTCGAAAACGAGGTCGCACGATTGAAAAAGTTGGTGGGTGAGggaggtggtggatcgggtggaggTTCTGATAAAGCGGAATTAATTCCCGAGCCTCCAAAAAAGAAGAGAAAGGTCGAATTGGGTACCGCCGCCAATTTTAATCAAAGTGGGAATATTGCTGATAGCTCATTGTCCTTCACTTTCACTAACAGTAGCAACGGTCCGAACCTCATAACAACTCAAACAAATTCTCAAGCGCTTTCACAACCAATTGCCTCCTCTAACGTTCATGATAACTTCATGAATAATGAAATCACGGCTAGTAAAATTGATGATGGTAATAATTCAAAACCACTGTCACCTGGTTGGACGGACCAAACTGCGTATAACGCGTTTGGAATCACTACAGGGATGTTTAATACCACTACAATGGATGATGTATATAACTATCTATTCGATGATGAAGATACCCCACCAAACCCAAAAAAAGAGtaa |
| eMagB-p65AD | atgggacataccctctacgcgccggggggttatgacatcatgggttacctcagacagatcagaaaccggccgaacccacaagtggagctgggacccgtcgacctctcctgcgccctcgtgctgtgtgaccttaagcagaaggacacccctgtggtgtacgcctccgaagcattcctggagatgaccgggtacaacagacacgaagtgctgggacggaactgccgcttcctgcaatccccggatggaatggtgaagcctaagtcaacccgcaaatacgtggactccaacactatctacaccatgaagaaggccattgaccgcaatgctgaggtgcaagtggaagtggtgaacttcaagaagaacggacagcgcttcgtcaacttcctgactatgattcccgtgcg |

| | |
|---|---|
| | ggacgaaaccggcgaataccggtacagcatcgggtttcagtgcgagactgagGGCGGTGGC<br>GGCAGTGGTGGAATGAAGCAACTCGAGGACAAGGTTGAGGAACTGC<br>TGAGTAAGAATTACCACCTCGAAAACGAGGTCGCACGATTGAAAAAGT<br>TGGTGGGTGAGggaggtggtggatcgggtggaggttctgaattccaataccttccagacacgg<br>atgatcgccatcgaatcgaagagaagagaaaacgcacctacgaaacgttcaaatctattatgaaga<br>aatctcccttcagtgggccgacggacccaaggccgccaccgcgaaggatagccgttccatcaagaa<br>gctcagcttctgtacccaaacccgccccacagccttacccctttacttcctccctctccactatcaactacg<br>atgagttccccacgatggtttttccttcaggacagatatcccaagcgagcgccctcgcaccagccccac<br>cacaagtgcttcctcaggcccctgcgcctgctccggcaccggcgatggtgagcgctctggctcaagct<br>cccgcgccagtccctgttttggcaccagggccacctcaggcagtggctccgcccgctccaaaacctac<br>tcaagcgggcgaaggaactctgagcgaggcgctcctgcagcttcaatttgatgacgaagatctcggc<br>gcactcctcggtaattcaacggaccccgctgttttcactgacctggcaagcgtggataactctgaattcc<br>aacagctccttaaccagggcataccggtcgcgcctcatacaactgaaccaatgctgatggaatatccg<br>gaggcaataaccagacttgtgacgggggcgcagcgaccgcctgatccagcacccgcaccgcttgg<br>ggcgcctggcttgcccaatggactcctttctggcgacgaggacttttccagcatcgcagacatggactttt<br>ctgcactcctttctcagatttcctcataa |
| eMagB-Msn2AD | atgggacataccctctacgcgccggggggttatgacatcatgggttacctcagacagatcagaaacc<br>ggccgaacccacaagtggagctgggacccgtcgacctctcctgcgccctcgtgctgtgtgaccttaag<br>cagaaggacacccctgtggtgtacgcctccgaagcattcctggagatgaccgggtacaacagacac<br>gaagtgctgggacggaactgccgcttcctgcaatccccggatggaatggtgaagcctaagtcaaccc<br>gcaaatacgtggactccaacactatctacaccatgaagaaggccattgaccgcaatgctgaggtgca<br>agtggaagtggtgaacttcaagaagaacggacagcgcttcgtcaacttcctgactatgattcccgtgcg<br>ggacgaaaccggcgaataccggtacagcatcgggtttcagtgcgagactgagGGCGGTGGC<br>GGCAGTGGTGGAATGAAGCAACTCGAGGACAAGGTTGAGGAACTGC<br>TGAGTAAGAATTACCACCTCGAAAACGAGGTCGCACGATTGAAAAAGT<br>TGGTGGGTGAGggaggtggtggatcgggtggaggttctggccctaaaaagaagcgtaaagtc<br>ACGGTCGACCATGATTTCAATAGCGAAGATATTTTATTCCCCATAGAAA<br>GCATGAGTAGTATACAATACGTGGAGAATAATAACCCAAATAATATTAAC<br>AACGATGTTATCCCGTATTCTCTAGATATCAAAAACACTGTCTTAGATAG<br>TGCGGATCTCAATGACATTCAAAATCAAGAAACTTCACTGAATTTGGG<br>GCTTCCTCCACTATCTTTCGACTCTCCACTGCCCGTAACGGAAACGAT<br>ACCATCCACTACCGATAACAGCTTGCATTTGAAAGCTGATAGCAACAAA<br>AATCGCGATGCAAGAACTATTGAAAATGATAGTGAAATTAAGAGTACTA<br>ATAATGCTAGTGGCTCTGGGGCAAATCAATACACAACTCTTACTTCACC<br>TTATCCTATGAACGACATTTTGTACAACATGAACAATCCGTTACAATCAC<br>CGTCACCTTCATCGGTACCTCAAAATCCGACTATAAATCCTCCCATAAA<br>TACAGCAAGTAACGAAACTAATTTATCGCCTCAAACTTCAAATGGTAAT<br>GAAACTCTTATATCTCCTCGAGCCCAACAACATACGTCCATTAAAGATA<br>ATCGTCTGTCCTTACCTAATGGTGCTAATTCGAATCTTTTCATTGACACT<br>AACCCAAACAATTTGAACGAAAAACTAAGAAATCAATTGAACTCAGATA<br>CAAATTCATATTCTAACTCCATTTCTAATTCAAACTCCAATTCTACGGGT<br>AATTTAAATTCCAGTTATTTTAATTCACTGAACATAGACTCCATGCTAGAT<br>GATTACGTTTCTAGTGATCTCTTATTGAATGATGATGATGATGACACTAA<br>TTTATCACGCCGAAGATTTAGCGACGTTATAACAAACtaa |
| eMagB-VP16AD | atgggacataccctctacgcgccggggggttatgacatcatgggttacctcagacagatcagaaacc<br>ggccgaacccacaagtggagctgggacccgtcgacctctcctgcgccctcgtgctgtgtgaccttaag<br>cagaaggacacccctgtggtgtacgcctccgaagcattcctggagatgaccgggtacaacagacac<br>gaagtgctgggacggaactgccgcttcctgcaatccccggatggaatggtgaagcctaagtcaaccc |

| | |
|---|---|
| | gcaaatacgtggactccaacactatctacaccatgaagaaggccattgaccgcaatgctgaggtgcaagtggaagtggtgaacttcaagaagaacggacagcgcttcgtcaacttcctgactatgattcccgtgcgggacgaaaccggcgaataccggtacagcatcgggtttcagtgcgagactgagGGCGGTGGCGGCAGTGGTGGAATGAAGCAACTCGAGGACAAGGTTGAGGAACTGCTGAGTAAGAATTACCACCTCGAAAACGAGGTCGCACGATTGAAAAAGTTGGTGGGTGAGggaggtggtggatcgggtggaggttctGGCCCTAAAAAGAAGCGTAAAGTCGCCCCCCCGACCGATGTCAGCCTGGGGGACGAGCTCCACTTAGACGGCGAGGACGTGGCGATGGCGCATGCCGACGCGCTAGACGATTTCGATCTGGACATGTTGGGGGACGGGGATTCCCCGGGTCCGGGATTTACCCCCCACGACTCCGCCCCCTACGGCGCTCTGGATATGGCCGACTTCGAGTTTGAGCAGATGTTTACCGATGCCCTTGGAATTGACGAGTACGGTGGGtaa |
| LexA-eMagAF | ATGccgaaaaagaaacgcaaagttggtagtAAAGCATTAACTGCTAGACAACAGGAGGTATTTGATTTGATTCGAGATCATATTTCTCAGACAGGGATGCCTCCAACCCGTGCCGAGATCGCCCAACGATTGGGATTTCGTAGTCCAAACGCCGCAGAAGAACACTTAAAGGCATTGGCCAGAAAGGGTGTCATAGAAATCGTGTCCGGTGCCAGTAGAGGGATTCGACTTTTACAGGAGGAAGAAGAGGGACTTCCTTTGGTAGGTAGAGTGGCTGCTGGGGAGCCATTATTAGCCCAACAACATATCGAAGGTCATTACCAAGTTGACCCAAGTCTTTTCAAGCCTAACGCAGACTTCTTGCTTAGAGTCAGTGGGATGTCAATGAAGGACATAGGTATTATGGACGGAGACTTGTTGGCTGTTCACAAAACACAGGATGTCAGAAACGGACAGGTGGTAGTCGCACGAATTGACGATGAAGTCACTGTTAAACGATTGAAAAAGCAGGGTAATAAAGTGGAGTTGTTACCTGAAAATTCTGAGTTCAAACCAATTGTTGTGGACCTTAGACAACAGTCATTTACAATCGAGGGTTTGGCAGTCGGTGTCATCAGAAACGGTGATTGGTTGggttctggaggcggaggctccggtggtggtggaagtggcggtggcggatccatgggacacactctttacgcccctggaggatacgacattatgggatatttggatcagattgcgaaccgcccaaaccctcaggtcgaactggggcctgtggacctgtcatgtgccctgatcctgtgcgatctgaagcaaaaggacactccgatcgtctacgcctcggaagccttcttggagatgaccggatacaacagacatgaggtgctcggcaggaactgcagattcctgcagtcccccgacgggatggtgaaaccaaagtcgactcgcaaatatgtggactcgaacacgatcttcaccatcaagaaggccatcgaccggaacgccgaggtccaggtggaggtggtcaactttaagaagaacggccagcggttcgtgaactttctgaccatcattccggtccgggatgaaaccggagagtacagatactccatcggattccagtgcgaaaccgaaTAA |
| NLS-VP16-EL222 | ATGGGCCCTAAAAAGAAGCGTAAAGTCGCCCCCCCGACCGATGTCAGCCTGGGGGACGAGCTCCACTTAGACGGCGAGGACGTGGCGATGGCGCATGCCGACGCGCTAGACGATTTCGATCTGGACATGTTGGGGGACGGGGATTCCCCGGGTCCGGGATTTACCCCCCACGACTCCGCCCCCTACGGCGCTCTGGATATGGCCGACTTCGAGTTTGAGCAGATGTTTACCGATGCCCTTGGAATTGACGAGTACGGTGGGGAATTCGGGGCAGACGACACACGCGTTGAGGTGCAACCGCCGGCGCAGTGGGTCCTCGACCTGATCGAGGCCAGCCCGATCGCATCGGTCGTGTCCGATCCGCGACTCGCCGACAATCCGCTGATCGCCATCAACCAGGCCTTCACCGACCTGACCGGCTATTCCGAAGAAGAATGCGTCGGCCGCAATTGCCGATTCCTGGCAGGTTCCGGCACCGAGCCGTGGCTGACCGACAAGATCCGCCAAGGCGTGCGCGAGCACAAGCCGGTGCTGGTCGAGATCCTGAACTACAAGAAGGACGGCACGCCGTTCCGCAATGCCGTGCTCGTTGCACCGATCTACGATGACGACGACGAGCTTCTCTATTTCCTCGGCAGCCAGGTCGAAGTCGACGACGACCAGCCCAACATGGGCATGGCGCGCCGCGAACGCGCCGCG |

| | |
|---|---|
| | GAAATGCTCAAGACGCTGTCGCCGCGCCAGCTCGAGGTTACGACGCT<br>GGTGGCATCGGGCTTGCGCAACAAGGAAGTGGCGGCCCGGCTCGG<br>CCTGTCGGAGAAAACCGTCAAGATGCACCGCGGGCTGGTGATGGAA<br>AAGCTCAACCTGAAGACCAGTGCCGATCTGGTGCGCATTGCCGTCGA<br>AGCCGGAATCTAA |
| NLS-VP16-EL22<br>2(A79Q) | ATGGGCCCTAAAAAGAAGCGTAAAGTCGCCCCCCCGACCGATGTCAG<br>CCTGGGGGACGAGCTCCACTTAGACGGCGAGGACGTGGCGATGGCG<br>CATGCCGACGCGCTAGACGATTTCGATCTGGACATGTTGGGGGACGG<br>GGATTCCCCGGGTCCGGGATTTACCCCCCACGACTCCGCCCCCTACG<br>GCGCTCTGGATATGGCCGACTTCGAGTTTGAGCAGATGTTTACCGATG<br>CCCTTGGAATTGACGAGTACGGTGGGGAATTCGGGGCAGACGACACA<br>CGCGTTGAGGTGCAACCGCCGGCGCAGTGGGTCCTCGACCTGATCG<br>AGGCCAGCCCGATCGCATCGGTCGTGTCCGATCCGCGACTCGCCGA<br>CAATCCGCTGATCGCCATCAACCAGGCCTTCACCGACCTGACCGGCT<br>ATTCCGAAGAAGAATGCGTCGGCCGCAATTGCCGATTCCTGCAAGGT<br>TCCGGCACCGAGCCGTGGCTGACCGACAAGATCCGCCAAGGCGTGC<br>GCGAGCACAAGCCGGTGCTGGTCGAGATCCTGAACTACAAGAAGGA<br>CGGCACGCCGTTCCGCAATGCCGTGCTCGTTGCACCGATCTACGATG<br>ACGACGACGAGCTTCTCTATTTCCTCGGCAGCCAGGTCGAAGTCGAC<br>GACGACCAGCCCAACATGGGCATGGCGCGCCGCGAACGCGCCGCG<br>GAAATGCTCAAGACGCTGTCGCCGCGCCAGCTCGAGGTTACGACGCT<br>GGTGGCATCGGGCTTGCGCAACAAGGAAGTGGCGGCCCGGCTCGG<br>CCTGTCGGAGAAAACCGTCAAGATGCACCGCGGGCTGGTGATGGAA<br>AAGCTCAACCTGAAGACCAGTGCCGATCTGGTGCGCATTGCCGTCGA<br>AGCCGGAATCTAA |
| NLS-VP16-EL22<br>2(AQTrip) | ATGGGCCCTAAAAAGAAGCGTAAAGTCGCCCCCCCGACCGATGTCAG<br>CCTGGGGGACGAGCTCCACTTAGACGGCGAGGACGTGGCGATGGCG<br>CATGCCGACGCGCTAGACGATTTCGATCTGGACATGTTGGGGGACGG<br>GGATTCCCCGGGTCCGGGATTTACCCCCCACGACTCCGCCCCCTACG<br>GCGCTCTGGATATGGCCGACTTCGAGTTTGAGCAGATGTTTACCGATG<br>CCCTTGGAATTGACGAGTACGGTGGGGAATTCGGGGCAGACGACACA<br>CGCGTTGAGGTGCAACCGCCGGCGCAGTGGGTCCTCGACCTGATCG<br>AGGCCAGCCCGATCGCATCGATTGTGTCCGATCCGCGACTCGCCGAC<br>AATCCGATTATCGCCATCAACCAGGCCTTCACCGACCTGACCGGCTAT<br>TCCGAAGAAGAATGCGTCGGCCGCAATTGCCGATTCCTGCAAGGTTC<br>CGGCACCGAGCCGTGGCTGACCGACAAGATCCGCCAAGGCGTGCGC<br>GAGCACAAGCCGGTGCTGGTCGAGATCCTGAACTACAAGAAGGACG<br>GCACGCCGTTCCGCAATGCCGTGCTCATTGCACCGATCTACGATGAC<br>GACGACGAGCTTCTCTATTTCCTCGGCAGCCAGGTCGAAGTCGACGA<br>CGACCAGCCCAACATGGGCATGGCGCGCCGCGAACGCGCCGCGGA<br>AATGCTCAAGACGCTGTCGCCGCGCCAGCTCGAGGTTACGACGCTG<br>GTGGCATCGGGCTTGCGCAACAAGGAAGTGGCGGCCCGGCTCGGCC<br>TGTCGGAGAAAACCGTCAAGATGCACCGCGGGCTGGTGATGGAAAA<br>GCTCAACCTGAAGACCAGTGCCGATCTGGTGCGCATTGCCGTCGAAG<br>CCGGAATCtaa |
